# Computational prediction and characterization of cell-type-specific and shared binding sites

**DOI:** 10.1101/2022.05.06.490975

**Authors:** Qinhu Zhang

**Affiliations:** Translational Medical Center for Stem Cell Therapy and Institute for Regenerative Medicine, Shanghai East Hospital, Bioinformatics Department, School of Life Sciences and Technology, Tongji University, Siping Road 1239, Shanghai 200092, China

## Abstract

Cell-type-specific gene expression is maintained in large part by transcription factors (TFs) selectively binding to distinct sets of sites in different cell types. Recent research works have provided evidence that such cell-type-specific binding is determined by TF’s intrinsic sequence preferences, cooperative interactions with cofactors, cell-type-specific chromatin landscapes, and 3D chromatin interactions. However, computational prediction and characterization of cell-type-specific and shared binding sites is rarely studied. In this paper, we propose two computational approaches for predicting and characterizing cell-type-specific and shared binding sites by integrating multiple types of features, in which one is based on XGBoost and another is based on convolutional neural network (CNN). To validate the performance of our proposed approaches, ChIP-seq datasets of 10 binding factors were collected from the GM12878 (lymphoblastoid) and K562 (erythroleukemic) human hematopoietic cell lines, each of which was further categorized into cell-type-specific (GM12878-specific and K562-specific) and shared binding sites. Then, multiple types of features for these binding sites were integrated to train the XGBoost-based and CNN-based models. Experimental results show that our proposed approaches significantly outperform other competing methods on three classification tasks. To explore the contribution of different features, we performed ablation experiments and feature importance analysis. Consistent with previous studies, we find that chromatin features are major contributors in which chromatin accessibility is the best predictor. Moreover, we identified independent feature contribution for cell-type-specific and shared sites through SHAP values, observing that chromatin features play a main role in the cell-type-specific sites while motif features play a main role in the shared sites. Beyond these observations, we explored the ability of the CNN-based model to predict cell-type-specific and shared binding sites by excluding or including DNase signals, showing that chromatin accessibility significantly improves the prediction performance. Besides, we investigated the generalization ability of our proposed approaches to different binding factors in the same cellular environment or to the same binding factors in the different cellular environments.

## INTRODUCTION

Transcription factors (TFs) play an integral role in the transcriptional regulatory networks by binding to their cognate motifs at specific locations to promote or repress the activities of other TFs, co-factors, chromatin modifiers, and the transcriptional machinery (1). However, the limited number of TFs encoded by the human genome (∼ 1600 (2)) is engaged in regulating the transcription activities of a large diversity of cell types, resulting in each individual TF being often reused in multiple distinct cell types and developmental stages. This observation raises the questions of how TFs choose their specific binding sites in distinct cell types and what genomic features can discriminate between cell-type-specific and shared binding sites (CSSBS). The process of discriminating these binding sites is difficult for most TFs, since they have very similar cognate motifs and sequence specificities.

Over the past several decades, the identification of transcription factor binding sites (TFBS) has made great progress, with both benefitting from the development of high-throughput sequencing technologies and the advancement of computational approaches. Specifically, ChIP-seq (3) provides an opportunity for viewing genome-wide interactions between DNA and specific TFs; Protein Binding Microarrays (PBMs) (4) have enabled large-scale characterization of TF-DNA binding in a high-throughput manner without considering the influence of cofactors on predicting TFBS. As a result, massive binding data provide an unprecedented opportunity to develop computational approaches to predict TFBS. MEME (5) discovered TF-DNA binding motifs by searching for repeated, ungapped sequence patterns that occur in the biological sequences. gkm-SVM (6) detected functional regulatory elements in DNA sequences by using gapped *k*-mer features and Support Vector Machine (SVM). Recently, Deep Learning (DL) has achieved amazing performance in many fields, inspiring researchers to develop DL-based methods for predicting TFBS (7-16). Typically, DeepBind (7) and DeepSea (8) applied Convolutional Neural Network (CNN) to accurately predict diverse molecular phenotypes, including TF binding from DNA sequences. DanQ (9) predicted TF-DNA binding motifs and prioritized functional SNPs by combining CNN with recurrent neural network (RNN). However, either traditional methods or DL-based methods focus on discriminating between binding and non-binding sites. This type of task (TFBS prediction) is relatively simple as the sequence specificities of binding and non-binding sites are often significantly different, so the methods using DNA sequences alone can achieve impressive performance. On the contrary, the task of accurately predicting CSSBS is more difficult and remains a lot of challenges.

The article (17) reviewed the various sequence and chromatin determinants of cell type-specific TF binding specificity and identified the current challenges and opportunities associated with computational approaches to characterize, impute, and predict cell type-specific TF binding patterns. As far as we know, computational approaches are rarely proposed to systematically study cell-type-specific and shared binding mechanisms. The most relevant research (18) analyzed hundreds of ChIP-seq datasets to explore the contributions of DNA sequence signal, histone modifications, and DNase accessibility to cell-type-specific binding, and proposed a discriminative framework for learning DNA sequences and chromatin signals that predict cell-type-specific TF binding. This work trained an SVM-based model that uses flexible *k*-mer patterns to capture DNA sequence signals more accurately than traditional motif approaches, and trained SVM spatial chromatin signatures to model local histone modifications and DNase accessibility, concluding that DNase accessibility can explain cell-type-specific binding for many factors. For better predicting cell-type-specific binding sites, several computational algorithms have been developed by combining DNA sequences with chromatin accessibility and histone modifications (19-21); however, they still focus on distinguishing binding sites from non-binding sites in distinct cell types. To study cell-type-specific and shared binding specificities, another relevant work (22) constructed an integrative approach to study estrogen receptor α (ER) through the integration of experimental measurements of TF binding, chromatin accessibility, DNA methylation, and mRNA levels, finding that ER exhibits two distinct modes for type-specific and shared binding sites; however, this study is limited to a few nuclear receptors and does not provide a computational approach for directly discriminating CSSBS.

In this paper, we present an XGBoost-based model for predicting and characterizing CSSBS through the integration of multiple types of features separately characterizing sequence specificities, cooperative interactions with cofactors, cell-type-specific chromatin landscapes, and 3D chromatin interactions. Previous studies have revealed that there is a strong correlation between cell-type-specific binding and sequence specificities, cofactors, chromatin landscapes (17). Additionally, chromatin conformation and DNA looping can also influence TF occupancy or enhancer selection in a cell-type-specific manner (23,24) which can be characterized by chromosome conformation capture assays such as Hi-C (25) providing high-resolution data on 3D chromatin interactions. In the light of the successful application of Deep Learning in many fields, we also present a CNN-based model for predicting CSSBS through the integration of DNA sequences and chromatin accessibility signals. Specifically, (i) we collected peaks and filtered bams of 10 different binding factors from the GM12878 and K562 cell lines, and then utilized DESeq2 (26) to differentially select GM12878-specific, K562-specific, and their shared binding sites; (ii) we applied MEME (5) to separately analyze the sequence preferences for these categorized binding sites, and ran LSGKM (27) to explain the difficulty of predicting CSSBS using DNA sequences alone; (iii) we downloaded chromatin landscape-related data, cofactor-related motifs and chromatin interaction-related data from public databases, which were then used to generate corresponding features for each site; (iv) on the basis of these generated features, we designed an XGBoost-based model to predict CSSBS, allowing us to better leverage multiple types of features to learn more discriminative patterns; (v) we also designed a CNN-based model to successfully predict CSSBS by encoding DNA sequences and chromatin accessibility signals together. Experimental results show that our proposed approaches can accurately predict CSSBS and significantly outperforms other competing methods. We conducted ablation experiments to explore the relative importance of different feature sets, finding that chromatin features are major contributors to predicting CSSBS. Furthermore, we performed an analysis of feature importance (SHAP values) for cell-type-specific and shared sites respectively to identify independent feature contributions, observing that chromatin features play a main role in the cell-type-specific sites while motif features play a main role in the shared sites. Beyond these observations, we also explored the ability of our proposed CNN-based method to predict CSSBS by excluding or including DNase signals, showing that chromatin accessibility can significantly improve the prediction performance. What’s more, we investigated the generalization ability of our proposed approaches to different binding factors in the same cellular environment or to the same binding factors in the different cellular environments, revealing that there exist some similar patterns of CSSBS between different binding factors in the same cellular environment but different patterns between the same binding factors in the different cellular environment.

## MATERIAL AND METHODS

### Data collection

ChIP-seq datasets of 10 binding factors (CEBPB, CTCF, FOS, JUNB, MAX, MYC, POLR2A, RAD21, SP1, and YY1) from the GM12878 and K562 cell lines were collected from the ENCODE Project (28), which meet the two criteria: (i) each dataset exists both in the GM12878 and K562 cell lines; (ii) each dataset has at least two replicates. Briefly, for each dataset in each cell line, the raw reads (fastq files) of all replicates were obtained from ENCODE, and then aligned to the hg19 reference genome keeping only uniquely aligned reads; these filtered alignments (bam files) were used to generate binding sites by using the peak calling software SPP (29) with default parameters. The standard data processing pipeline is available from the ENCODE DCC Github (https://github.com/ENCODE-DCC/chip-seq-pipeline2). In addition, we collected a cohort of publicly available chromatin landscapes including DNase-seq, 11 ChIP-seq histone modifications (H2A.Z, H3K4me1, H3K4me2, H3K4me3, H3K9ac, H3K9me3, H3K27ac, H3K27me3, H3K36me3, H3K79me2, and H4K20me1), MNase-seq, total RNA-seq, and DNA methylation from the Roadmap Epigenomics Project (30), jointly forming the chromatin environments of the GM12878 and K562 cell lines. We obtained 5kb resolution Hi-C contact maps of GM12878 and K562 from another work (31), capturing the contact information of all potential regulatory sites. All datasets used in this work are summarized in Supplementary tables 1, 2, and 3.

### Differential binding sites preparation

We defined differential binding sites as peaks that are significant differences between GM12878 and K562. Briefly, for each binding factor, (i) we merged all peaks from the two cell lines, and used corresponding filtered alignments (bam files) to count the number of reads from each cell line falling into each merged peak by utilizing bedtools (https://bedtools.readthedocs.io/en/latest/); (ii) optionally, we normalized the read counts by using the equation (1) to reduce the influence of sequencing depth and peak length; (iii) we ran DESeq2 (26) on the normalized read counts to compute the difference significance between GM12878 and K562; (iv) peaks with *q*-value less than 0.05 and log2 fold-change less than -2 were chosen as GM12878-specific binding sites, peaks with *q*-value less than 0.05 and log2 fold-change greater than 2 were chosen as K562-specific binding sites, and peaks with *q*-value greater than 0.1 and the absolute value of log2 fold-change less than 1 were chosen as shared binding sites, while the remaining peaks were abandoned. Note that we can control the number and quality of selected peaks by modifying the *q*-value or log2 fold-change.

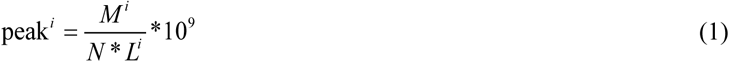

where *M, N*, and *L* represent the mapped reads of the *i*-th peak, the total mapped reads of all peaks, and the length of the *i*-th peak, respectively.

In this work, we applied a split-by-chromosome strategy to divide all selected peaks into training, test, and validation data, where chromosomes 8 and 16 were used as the test set, chromosome 18 was used as the validation set while the remaining chromosomes except Y were used as the training set. Moreover, each peak was trimmed to 600bp length.

### Multiple tasks

Since all peaks were differentially divided into GM12878-specific binding sites, K562-specific binding sites, and their shared binding sites, correspondingly we designed three classification tasks in this study, separately named task-A, task-B, and task-C. More precisely, task-A means discriminating between GM12878-specific and K562-specific binding sites; task-B means discriminating between cell-type-specific and shared binding sites; task-C means discriminating among GM12878-specific, K562-specific, and shared binding sites. Therefore, task-A and task-B are two-class classification tasks whose labels are represented by 0 and 1 while task-C is a three-class classification task whose labels are represented by 0, 1, and 2. Note that task-B is the primary task discussed in this paper.

For evaluating the performance of our proposed methods on task-A and task-B, the AUC (Area Under the Receiver Operating Characteristic Curve) and PRAUC (Area Under the Precision-Recall Curve) were adopted; whereas for evaluating the performance of our proposed methods on task-C, the balanced accuracy allowing for imbalanced data and macro f1-score suitable for multi-class problems were adopted.

### Feature generation

Previous research works (17,18) have shown that cell-type-specific binding is in large part determined by TF’s intrinsic sequence preferences, cooperative interactions with cofactors, cell-type-specific chromatin landscapes, and 3D chromatin interactions. Therefore, we considered four types of feature sets in this paper, including sequence features, motif features, chromatin features, and chromatin interaction features.

#### A. Sequence features

Considering that sequence-based features play an important role in identifying TFBS, we applied two different ways to generate sequence features. More precisely, (i) a more concise version of gapped k-mer frequency vectors (gkm-fvs) which significantly reduces the dimension of the original gkm-fvs were directly yielded by using the method (32) with default parameters, resulting in a total of 7680 features; or (ii) CNN-based sequence features were yielded from the outputs of the last convolutional layer in a CNN model pre-trained on the training set, resulting in a total of 500 features.

#### B. Motif features

Firstly, we ran MEME on each binding factor’s dataset to identify motifs that are directly relevant to their binding sites, and then collected physically-or functionally-interacting cofactors with high confidence scores (0.8 in this paper) through the protein-protein-interaction network (PPI) (33). Secondly, we merged all relevant factors, filtered those whose motifs do not exist in the JASPAR database (34), and acquired the corresponding positional weight matrices (PWMs)of the remaining factors. Finally, we scored DNA forward and reverse-complement sequences by the summation of the product of PWMs and sequence matrices encoded by one-hot, and then selected the top three scores as the motif features. Since each binding factor has a different number of relevant factors, its motif features have a total of *n*\*3 where *n* denotes the number of relevant factors.

#### C. Chromatin features

15 types of chromatin landscapes across the GM12878 and K562 cell lines were used in this paper. For each chromatin landscape, we calculated the minimum, mean, and maximum signal values of each peak (600bp length) representing the overall distribution of signals around the peak. Furthermore, we computed the difference of each chromatin landscape between GM12878 and K562 in terms of the minimum, mean, and maximum signal values. Therefore, a total of 135 (2*15*3+15*3) chromatin features were generated.

#### D. Chromatin interaction features

Hi-C sequencing technology quantifies contacts between all possible pairs of genomic loci making it possible to detect loops at kilobase resolution, thereby Hi-C contact maps of GM12878 and K562 at 5kb resolution were downloaded from public databases. Briefly, we first selected the top 20 regions interacting with each peak by the normalized contact counts, and then recorded the counts and distances between the peak and its top 20 interacting regions; we calculated the minimum, mean, and maximum values of the counts and distances, and computed the difference of these values between GM12878 and K562, resulting in a total of 18 (2*6+6) chromatin interaction features.

At last, the four types of feature sets were concatenated into vectors as the inputs to the XGBoost-based model described in the next section.

### Models construction

In this work, we constructed two models for computational prediction and characterization of CSSBS. One is based on Ensemble Learning and another is based on Deep Learning.

#### A. XGBoost classifier

XGBoost (35) is an optimized distributed gradient boosting library designed to be highly efficient, flexible, and portable, which implements machine learning algorithms under the Gradient Boosting framework for classification and regression tasks. In an XGBoost model, boosted trees are added into the model by optimizing the loss function from previous trees. For each binding dataset, we trained an XGBoost model using the four types of feature sets described above, and employed a grid-search strategy to select the best parameters from a fixed set of parameters (e.g, ‘max depth’: {5, 6, 7, 8}; ‘learning rate’: {0.001, 0.1, 0.3, 0.5, 1}); furthermore, we set the maximum number of iterations to 1000 and applied a validation-based early stopping strategy; in the end, the final model for this dataset was re-trained using the best parameters.

#### B. CNN classifier

Since CNN is suitable for directly dealing with DNA sequences, each peak is transformed into an *L* × 4 matrix where *L* denotes the length of this peak with one-hot format (A = [1, 0, 0, 0], C= [0, 1, 0, 0], G = [0, 0, 1, 0] and T = [0, 0, 0, 1]). For accurately predicting CSSBS, we integrated DNase signals of GM12878 and K562, the difference of DNase signals between GM12878 and K562, and the sequence matrix, jointly forming an *L* × 7 matrix. The CNN-based model is composed of three convolutional blocks in which each block is made up of a convolutional layer, a ReLU layer, a max-pooling layer as well as a dropout layer, and an output block which is made up of a fully-connected layer followed by a ReLU layer, a dropout layer, and a fully-connected layer followed by a sigmoid layer (for task-A and task-B) or a softmax layer (for task-C). Generally, the calculation process of the CNN-based model can be described by equations (2) and (3).

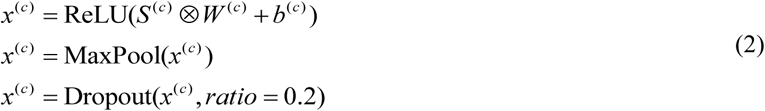

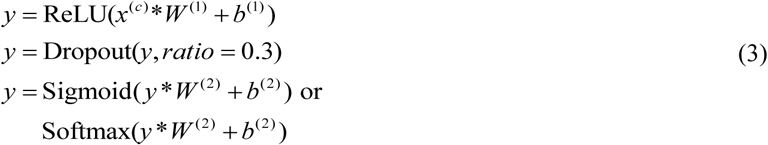

where *S*^(*c*)^, *x*^(*c*)^, *W*^(*c*)^, and *b*^(*c*)^ are input, output, weight matrix, and bias vector of the *c*^th^ convolutional block, respectively; ⨂ denotes the convolutional operation; *W*^(1)^ and *b*^(1)^, *W*^(2)^ and *b*^(2)^ denote the weights and biases of the two fully-connected layers.

Allowing for the issues that the initialization of weights and the selection of hyperparameters may influence the overall performance of the CNN-based model, we repeatedly warmed up the CNN classifier by randomly initializing weights and searching for the best hyperparameters from a fixed set (e.g. ‘learning rate’: {0.01, 0.001, 0.0001}). Then, we re-trained the best-performing CNN model which stores the best initialization values and hyper-parameters using the training set. The cross-entropy loss was used to train the CNN model, and optimized by the Adam optimization algorithm with a batch size of 300. All details of setting parameters can refer to our released code.

### Baseline methods

As far as we know, computational approaches for predicting cell-type-specific and shared sites are rarely proposed. The most relevant work (18) proposed an SVM-based discriminative framework for learning DNA sequence and chromatin signals that predict cell-type-specific TF binding, but it still focused on classifying binding and non-binding sites. Another relevant work (22) constructed an integrative approach to reveal distinct properties of cell-type-specific and shared transcription factor binding sites; however, this study did not provide a computational approach for directly discriminating CSSBS. In order to evaluate the performance of our proposed methods, we chose several kinds of representative methods as the baselines, which are widely used in the task of cell-type-specific TFBS prediction.

#### LSGKM

is an improved version of gkm-SVM for large-scale datasets which provides further advanced gapped *k*-mer-based kernel functions, resulting in considerably higher accuracy than gkm-SVM. Moreover, since gapped *k*-mer features have been proved to be better than *k*-mer features (6), we did use gapped *k*-mer features instead of *k*-mer features in this study.

#### CNN

is the designed CNN-based model excluding chromatin accessibility (DNase signals)in this study (see **Models construction**).

#### SVC

is an SVM-based method capable of performing binary and multi-class classification, which chooses ‘RBF’ as the kernel function for learning nonlinearity and uses the ‘one-vs-one’ decision function for multi-class classification.

#### Random Forest (RF)

is a typical method of Ensemble Learning which is composed of a diverse set of randomized decision trees, where each tree is built from a sample drawn with replacement from the training set with a random subset of features.

LSGKM and CNN used DNA sequences alone while SVC and RF took all types of features described above as input. Both SVC and RF were implemented using scikit-learn (36).

## RESULTS

### Overview of our proposed approaches for predicting cell-type-specific and shared binding sites

As we know, a great diversity of algorithms for TFBS prediction have achieved impressive performance even using DNA sequences alone, but they mainly focus on distinguishing binding sites from non-binding sites and rarely pay attention to predicting CSSBS. Furthermore, the task of discriminating CSSBS is thought to be more difficult than the task of TFBS prediction, as CSSBS may have similar sequence specificities while binding and non-binding sites show significantly-different sequence specificities. To accomplish the task of discriminating CSSBS, we first collected ChIP-seq datasets of 10 binding factors, including transcription factor, cohesion, cofactor, and RNA polymerase complex, from the GM12878 and K562 cell lines, and selected cell-type-specific and shared binding peaks by utilizing DESeq2and then developed an XGBoost-based method and a CNN-based method to predict and characterize these differential binding peaks. In the XGBoost-based model (Figure 1A), four types of feature sets including sequence features, motif features, chromatin features, and chromatin interaction features were first generated, and then XGBoost took these features as input and output the probabilities of classes. In the CNN-based model (Figure 1B), one-hot vectors derived from DNA sequences and DNase accessibility signals were first encoded together, and then CNN took these features as input and output the probabilities of classes. The details of our proposed approaches can refer to the ‘**Models construction**’ section and our released code. For the sake of simplicity, our proposed methods are abbreviated as XGBoost and CNN-plus respectively in the subsequent sections.

**Figure 1.**
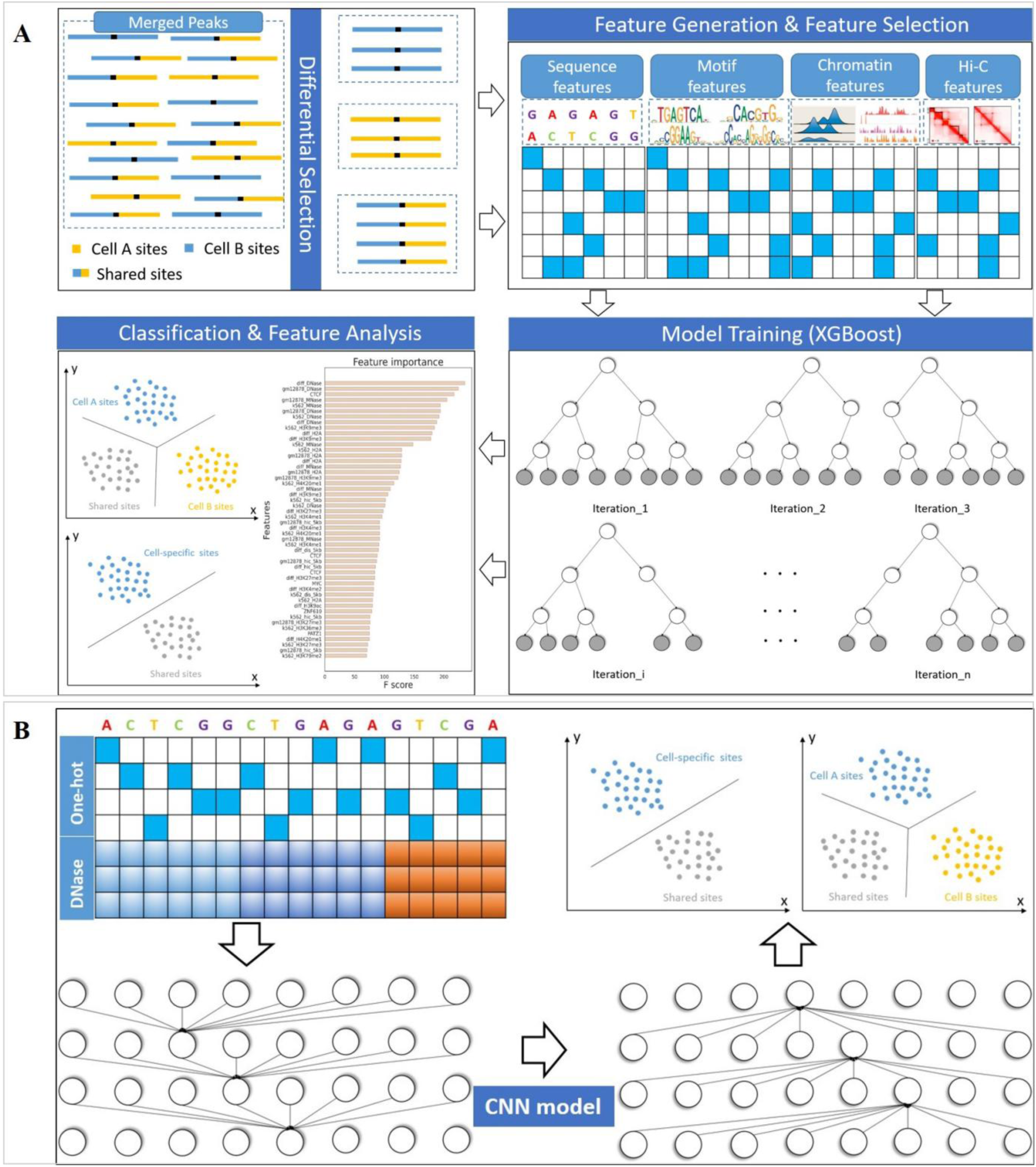
Schematic overview of our proposed approaches. (A) After selecting differential binding sites, the XGBoost-based model takes sequence features, motif features, chromatin features, and chromatin interaction features as inputs and outputs the probabilities of labels. (B) After selecting differential binding sites, the CNN-based model takes DNA sequences and chromatin accessibility signals as inputs and outputs the probabilities of labels. The two models are used to discriminate between GM12878-specific and K562-specific binding sites (task-A), discriminate between cell-type-specific and shared binding sites (task-B), and discriminate among GM12878-specific, K562-specific, and shared binding sites (task-C).

Figure 2A displays the distribution of binding peaks for CTCF and POLR2A (other binding factors can be found in Supplementary figure 1), in which GM12878-specific, K562-specific, shared, and abandoned peaks are labelled by different colors, implying that we can control the quality and number of selected peaks by adjusting the *q*-value and log2 fold-change. To validate the reasonability of these peaks, for CTCF, we extracted the DNase signals of GM12878 and K562 around each peak and plotted the heat maps of them. As shown in Figure 2B, the DNase signals of GM12878 are enriched in the GM12878-specific peaks and the DNase signals of K562 are enriched in the K562-specific peaks, while the DNase signals of GM12878 and K562 are both enriched in the shared peaks. For POLR2A, since it is the largest subunit of RNA polymerase II responsible for transcriptional activities, we searched for the nearest gene to each peak and extracted its corresponding gene expression level from RNA-seq. As shown in Figure 2C, in general, the genes for cell-type-specific peaks are differentially expressed (distributed on both sides) while the genes for shared peaks are similarly expressed (clustered on the diagonal line). Note that not all peaks of POLR2A are involved in synthesizing messenger RNA (mRNA) in the process of gene transcription, for example, enhances can also be transcribed into eRNA (37), so a few points are crossly distributed. Besides, we computed the Pearson correlation of DNase signals between GM12878 and K562 for cell-type-specific and shared binding peaks from 10 binding factors. As shown in Figure 2D, the Pearson correlation of shared binding peaks is almost all higher than that of cell-type-specific binding peaks. These observations demonstrate that the way of selecting cell-type-specific and shared binding peaks is reasonable and feasible.

**Figure 2.**
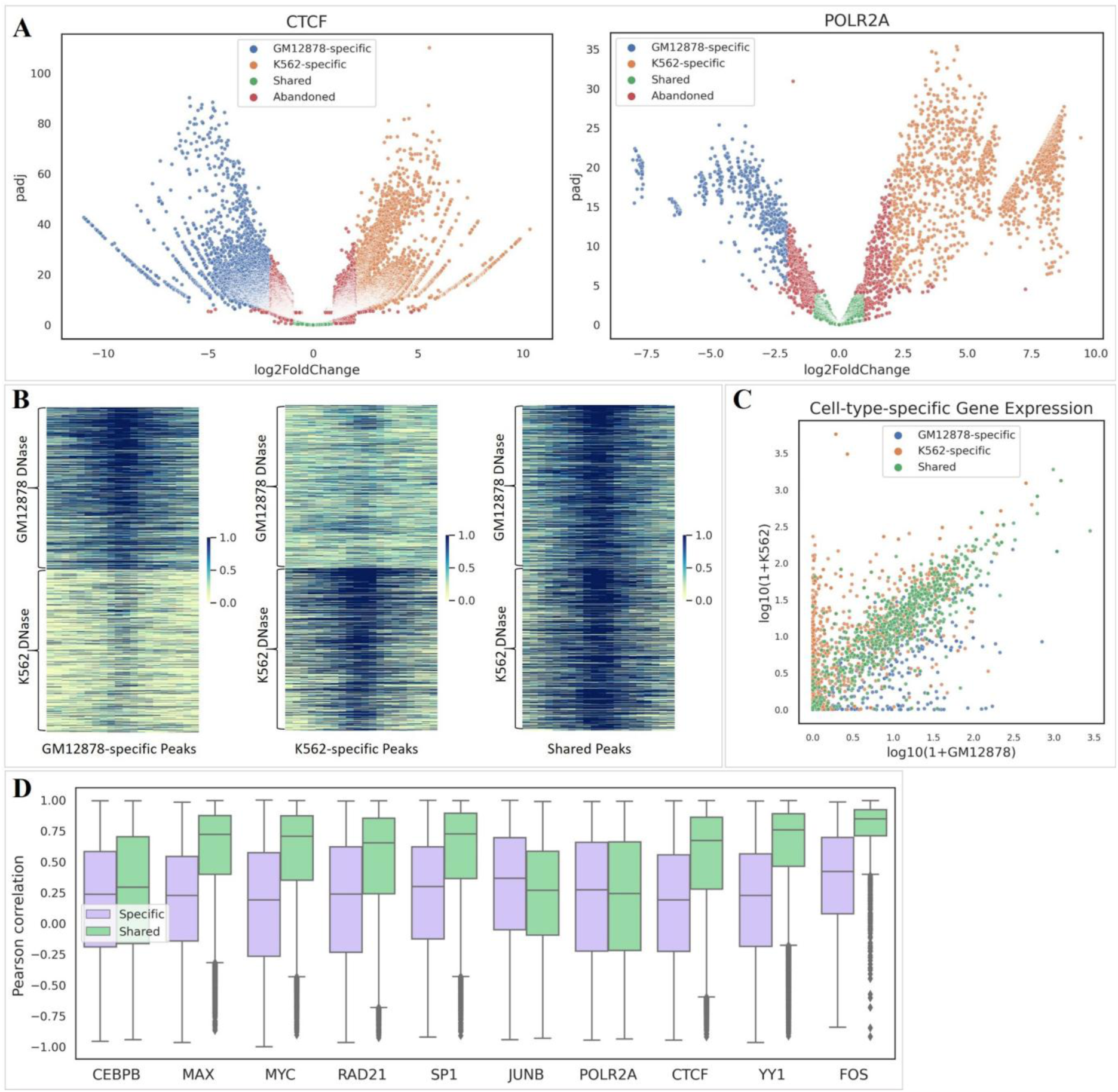
An analysis of cell-type-specific and shared binding peaks. (A) The distribution of binding peaks for CTCF and POLR2A where those with *q*-value less than 0.05 and log2 fold-change less than -2 are viewed as GM12878-specific binding sites, those with *q*-value less than 0.05 and log2 fold-change greater than 2 are viewed as K562-specific binding sites, and those with *q*-value greater than 0.1 and absolute value of the log2 fold-change less than 1 are viewed as shared binding sites, while those labelled by red color represent abandoned peaks. Note that we can control the number and quality of selected peaks by modifying the *q*-value and the log2 fold-change. (B) The visualization of CTCF DNase signals between GM12878 and K562 for GM12878-specific, K562-specific, and shared binding peaks. (C) The cell-type-specific gene expression (RNA-seq) of genes closest to POLR2A binding peaks between GM12878 and K562. (D) The Pearson correlation of DNase signals between cell-type-specific and shared binding peaks across 10 binding factors.

### The difficulty of predicting cell-type-specific and shared binding sites using DNA sequences alone

To demonstrate the difficulty of predicting CSSBS, we conducted a preliminary analysis of motif similarity by running MEME on GM12878-specific, K562-specific, and shared binding peaks respectively. As shown in Figure 3A (CEBPB and CTCF), the corresponding canonical motifs are all found in the GM12878-specific, K562-specific, and shared peaks respectively, and most of the identified cobinding factors are also overlapped. Furthermore, we used FIMO (38) to compare motif hits among GM12878-specific, K562-specific, shared, and non-binding peaks, finding that the ratio of motif hits to the total number of peaks and matched *p*-value are almost identical across the GM12878-specific, K562-specific, and shared peaks but higher than that of non-binding peaks (Figures 3B and 3C). These observations imply that the task of predicting CSSBS is more difficult than the task of predicting TFBS.

**Figure 3.**
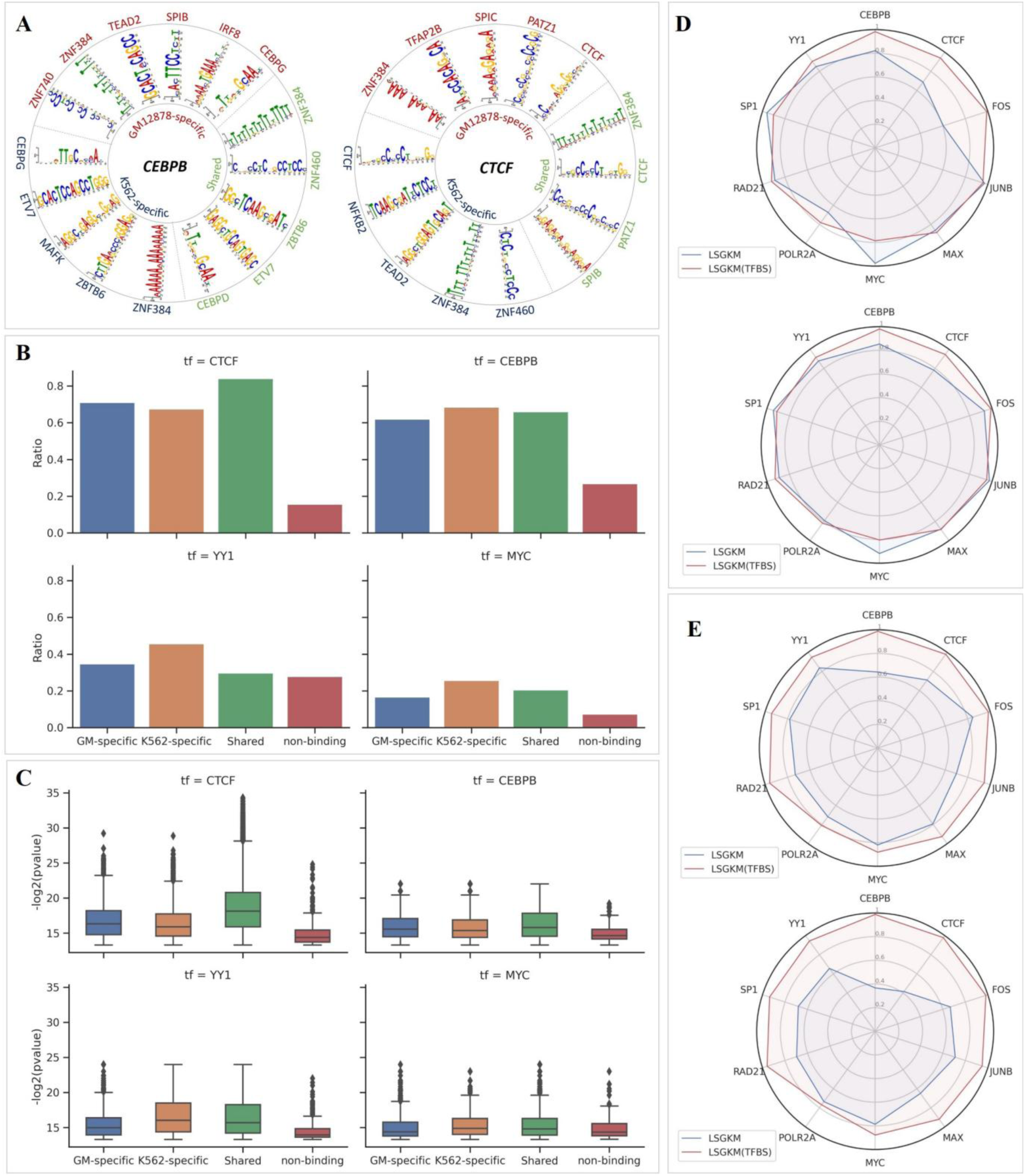
An analysis of the difficulty of predicting CSSBS using DNA sequences alone. (A) The identified motifs of GM12878-specific, K562-specific, and shared binding peaks by MEME for CEBPB and CTCF. (B) The ratio of found motif instances to all peaks across CTCF, CEBPB, YY1, and MYC. (C) The distribution of the –log2(p-value) of found motif instances across CTCF, CEBPB, YY1, and MYC. (D) The AUC and PRAUC comparison of LSGKM and LSGKM(TFBS) across all binding factors where ‘LSGKM’ means discriminating between GM12878-specific and K562-specific binding sites while ‘LSGKM(TFBS)’ means discriminating between binding sites (GM12878-specific and K562-specific binding peaks) and corresponding non-binding sites. (E) The AUC and PRAUC comparison of LSGKM and LSGKM(TFBS) across all binding factors where ‘LSGKM’ means discriminating between cell-type-specific and shared binding sites while ‘LSGKM(TFBS)’ means discriminating between binding sites (cell-type-specific and shared binding peaks) and corresponding non-binding sites.

As we mentioned above, the task of TFBS prediction is to distinguish binding sites from non-binding sites, achieving significant performance even using DNA sequences alone. To quantitatively analyze the difficulty of predicting CSSBS, we compare task-A and task-B with the task of TFBS prediction (see ‘**Multiple tasks**’). Briefly, for task-A, we performed LSGKM using DNA sequences derived from the GM12878-specific and K562-specific binding peaks; for task-B, we similarly performed LSGKM using DNA sequences derived from the cell-type-specific and shared binding peaks. However, the peaks for task-A or task-B were regarded as the positive set while the upstream regions of these peaks were extracted as the negative set in the task of TFBS prediction; then, we ran LSGKM on the positive and negative sets. As shown in Figures 3D and 3E, we find that (i) the overall performance of TFBS prediction is significantly better than that of task-A (the average AUC: 0.901 vs. 0.885; the average PRAUC: 0.891 vs. 0.83) and task-B (the average AUC: 0.936 vs. 0.758; the average PRAUC: 0.93 vs. 0.637), demonstrating that task-B (CSSBS prediction) is much more difficult than the task of TFBS prediction, and (ii) the performance of task-A is better than that of task-B (the average AUC: 0. 885 vs. 0.758; the average PRAUC: 0. 83 vs. 0.637), inferring that the sequence specificities between cell-type-specific binding sites are more discriminative than that between cell-type-specific and shared binding sites. Similarly, we can observe the same results when using CNN to do TFBS prediction (Supplementary figure 2).

### Chromatin features are major contributors to predicting cell-type-specific and shared binding sites

Since cell-type-specific binding is in large part determined by TF’s intrinsic sequence preferences, cooperative interactions with cofactors, cell-type-specific chromatin landscapes, and 3D chromatin interactions, four types of feature sets were generated as the inputs to XGBoost. We used the gkm-fvs or outputs of CNN to represent TF’s intrinsic sequence preferences, the motif scores from PPI to represent cooperative interactions with cofactors, 15 chromatin-related data (e.g, chromatin accessibility, histone modification, DNA methylation) to represent cell-type-specific chromatin landscapes, and the Hi-C contact maps to represent 3D chromatin interactions. To evaluate the relative importance of each feature set to the prediction performance of XGBoost, we conducted a series of ablation experiments by excluding one type of feature set from all sets. At first, we compared the effect of the gkm-fvs and CNN-based sequence features, finding that the performance of using CNN-based sequence features is better than that of using gkm-fvs (Figure 4A). We think that the high dimensionality of gkm-fvs is the main reason to be unfriendly to XGBoost. Therefore, we used CNN-based sequence features in the subsequent ablation experiments. As shown in Figure 4A, the performance of XGBoost without using sequence features is almost identical to the one using all features, meaning that sequence features have little effect on the performance of CSSBS prediction; the performance of XGBoost without using motif features or chromatin interaction features is slightly decreased while the performance of XGBoost without using chromatin features is significantly decreased, obviously indicating that chromatin features are major contributors to predicting CSSBS. These observations are also consistent with the findings in recent related works (19,20). Given the above observations, we remained motif features, chromatin features, and chromatin interaction features for XGBoost in the subsequent analysis or experiments.

**Figure 4.**
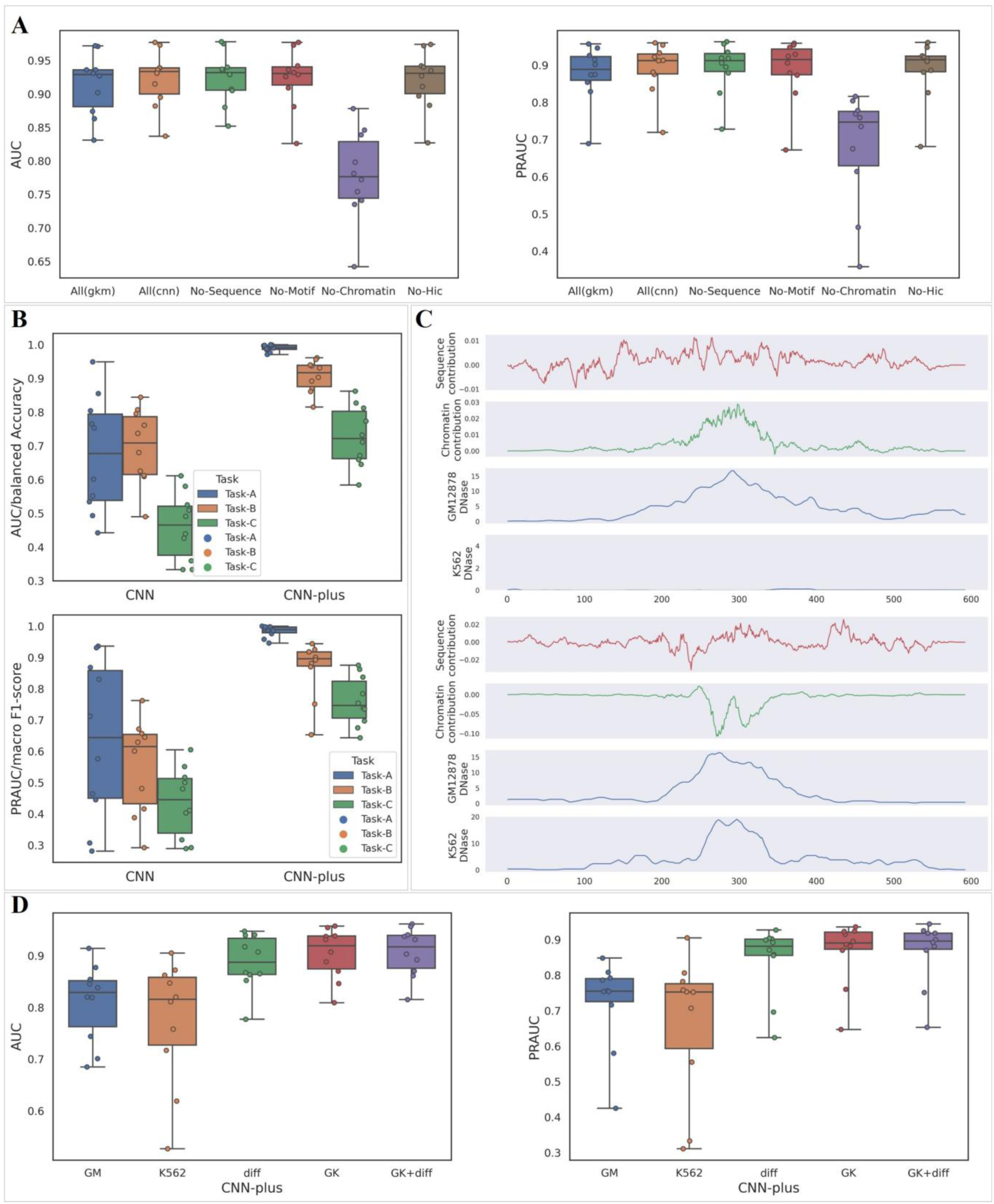
(A) The performance comparison of XGBoost on predicting CSSBS under different combinations of feature sets. (B) The performance comparison of CNN and CNN-plus on the task-A, task-B, and task-C. (C) The nucleotide-resolution contributions of DNA sequences and chromatin accessibility coupled with cell-type-specific DNase signals in which the top panel corresponds to a cell-type-specific binding site and the bottom panel corresponds to a shared binding site. (D) The performance comparison of CNN-plus on predicting CSSBS under different combinations of DNase-related features where diff denotes the difference of DNase signals between GM12878 and K562, and GK represents a combination of GM12878 and K562 DNase signals.

### Chromatin accessibility enhances CNN’s ability to predict cell-type-specific and shared binding sites

Without the cell-type-specific information on chromatin accessibility, those DL-based models that rely only on DNA sequences cannot distinguish the diverse TF binding profiles across different cell types. Recent methods, such as FactorNet (21) and Leopard (39), address this problem by considering both DNA sequence and chromatin accessibility, greatly improving the prediction performance. However, they still focus on the task of discriminating between binding and non-binding sites. On the contrary, we used GM12878-specific DNase signals, K562-specific DNase signals, and the difference between them as the cell-type-specific information on chromatin accessibility, together with DNA sequences to predict CSSBS. To verify the impact of the cell-type-specific chromatin accessibility on predicting CSSBS, we compared CNN with CNN-plus on the three tasks (task-A, task-B, and task-C), where CNN-plus takes DNA sequences and cell-type-specific chromatin accessibility signals as input while CNN uses DNA sequences alone. As shown in Figure 4B, we observe that CNN-plus significantly outperforms CNN on the three tasks, and the average performance on the three tasks is improved by about 33%, 26%, and 30% respectively, strongly supporting that chromatin accessibility can enhance CNN’s ability to predict CSSBS. Moreover, we computed the contribution of each base (nucleotide) to the prediction by applying DeepLIFT (40) which is a method for decomposing the output prediction of a neural network on a specific input by backpropagating the contributions of all neurons in the network to every feature of the input. This method assigns contribution scores according to the difference between the activation of each neuron to its ‘reference’ activation. Therefore, to investigate the relative contributions of DNA sequences and chromatin accessibility, we designed two ‘references’ for each input where one was equal to the input whose sequence-related values were set to 0 while another was equal to the input whose chromatin-related values were set to 0. In Figure 4C, the top panel shows an example of cell-type-specific binding peaks, and the bottom panel shows an example of shared binding peaks, observing that (i) the contribution of chromatin accessibility is more enriched than the one of DNA sequences and consistent with its cell-type-specific chromatin accessibility; (ii) cell-type-specific chromatin accessibility makes a positive contribution while shared chromatin accessibility makes a negative contribution, thereby providing more discriminative features for predicting CSSBS. The above observations demonstrate that the cell-type-specific information on chromatin accessibility is very important for such cell-type-specific predictive tasks.

To further explore the effect of each type of chromatin accessibility, we conducted some relevant ablation experiments by excluding one of three chromatin DNase signals. As shown in Figure 4D, we find that (i) CNN-plus using GM12878 or K562 DNase signals alone fails to predict CSSBS; (ii) CNN-plus using the difference of DNase signals between GM12878 and K562 can achieve very impressive performance, and the average performance is improved by about 8% (AUC) and 12% (PRAUC) compared with (i); (iii) CNN-plus using all chromatin DNase signals performs the best in all experiments. Overall, a combination of the three types of chromatin accessibility is essential for accurately predicting CSSBS.

### The overall performance of XGBoost on the three tasks

For XGBoost, we constructed three types of feature sets including motif features, chromatin features, and chromatin interaction features derived from PPI, chromatin landscapes, and Hi-C contact maps. To test the prediction performance of XGBoost, we compared it with LSGKM, CNN, SVC, Random Forest (RF), and CNN-plus on the three tasks (task-A, task-B, and task-C). As shown in Figures 5A (task-A) and 4B (task-B), LSGKM achieves very high predictive accuracy and performs much better than CNN even using DNA sequences alone, implying that gapped k-mer encoding is a better representative of sequence specificities between cell-type-specific binding sites than nucleotide-independent one-hot encoding. Moreover, XGBoost performs much better than LSGKM and CNN, demonstrating that three types of feature sets are better than sequence features; XGBoost outperforms SVC and RF, indicating that these features are better suitable for XGBoost; XGBoost is superior to CNN-plus, possibly benefitting from more classification features. For the multi-class classification (task-C), we get similar conclusions except that CNN-plus performs worse than SVC and RF, suggesting that the features learned by CNN-plus are more suitable for two-class classification. Compared with LSGKM, CNN, SVC, RF, and CNN-plus (Supplementary figure 3), for task-A, the average AUC of XGBoost is improved by about 11%, 32%, 4%, 3%, and 0.2% respectively, and the average PRAUC of XGBoost is improved by about 16%, 35%, 6%, 4%, and 0.4% respectively; for task-B, the average AUC of XGBoost is improved by about 17%, 23%, 12%, 9%, and 2% respectively, and the average PRAUC of XGBoost is improved by about 26%, 34%, 20%, 16%, and 3% respectively; for task-C, the average balanced accuracy of XGBoost is improved by about 33%, 3%, 2%, and 7% respectively, and the average macro F1-score of XGBoost is improved by about 38%, 2%, 1%, and 6% respectively. These performance gains show us that XGBoost can identify discriminative features from all feature sets and learn to accurately implement classification on the three tasks.

**Figure 5.**
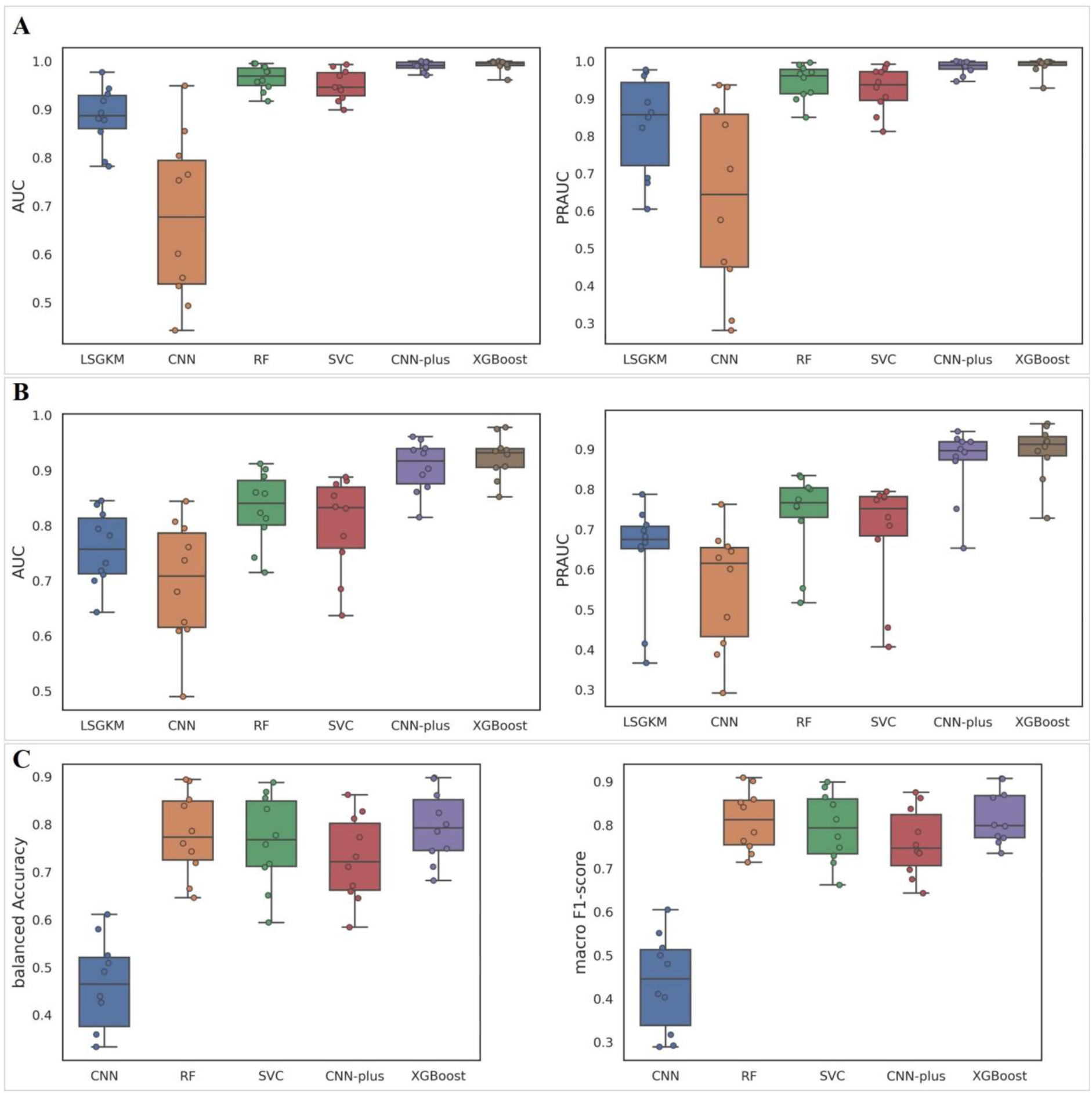
The overall performance of our proposed approaches. (A) The performance comparison of all methods on the task-A under the AUC and PRAUC metrics. (B) The performance comparison of all methods on the task-B under the AUC and PRAUC metrics. (C) The performance comparison of all methods on the task-C under the balanced Accuracy and macro F1-score metrics. The description of the three tasks please refer to the ‘**Multiple tasks**’ section.

From the above results, we find that XGBoost is slightly better than CNN-plus on the two-class classification tasks (task-A and task-B); however, XGBoost requires much more effort to do feature engineering while CNN-plus only needs DNA sequences and cell-type-specific chromatin accessibility, suggesting the potential advantage of applying CNN-plus to predict CSSBS. For the three-class classification task, RF can be used as an alternative to XGBoost but also needs much effort to do feature engineering.

### Feature importance analysis to identify independent feature contributions

The importance value of each feature was calculated by counting the number of its occurrences in all boosted trees. If a feature occurs more frequently in the nodes of multiple trees, it is more important. Through feature importance, we can further analyze the independent contribution of each feature instead of just analyzing the contribution of each feature set. Since we were mainly concerned about features of high importance, we selected the top 30 independent features to display. As shown in Figure 6A, we find that the most prominent feature is the ‘diff_DNase’ that denotes the difference of DNase signals between GM12878 and K562, demonstrating that this feature is the most important effector to predict CSSBS which is consistent with the finding in the analysis of CNN-plus, and that chromatin-related features account for the largest number of important factors among the top 30 independent features, again proving that chromatin features are major contributors to predicting CSSBS. In addition, for CTCF, its canonical motif feature is a very important effector ranked third since it has a long and strongly-specific motif; CTCF can act as a boundary factor often involved in forming chromatin loops (41,42), thus chromatin interaction features (k562_hic_5kb and gm12878_hic_5kb) play an important role. For JUNB (AP-1 transcription factor subunit), some cofactors physically-or functionally-interacting with it are the main factors where FOSL1 is often combined with JUNB to form JUN-FOS heterodimers which strongly binds to the TPA-response element (43); however, the top 30 features do not contain JUNB’s cognate motifs, which implies that JUNB’s motif features are not specific between cell-type-specific and shared binding sites. For POLR2A (RNA polymerase II subunit), except DNase features, RNA-seq features are the most important factors which represent their specific gene expression; the only motif feature among the top 30 features corresponds to TBP, which is a general transcription factor that functions at the core of the DNA-binding multiprotein factor TFIID, playing a role in the activation of eukaryotic genes transcribed by RNA polymerase II; chromatin interaction features are highly ranked since Hi-C experiments have proved that genome exists a large number of promoter-mediated chromatin loops such as promoter-promoter loops and enhancer-promoter loops. For RAD21 (cohesin complex subunit), it has been annotated as a factor without sequence specificity but can be combined with CTCF to form CTCF-mediated chromatin loops (41) or associated with MYC to control cohesin positioning and genome organization (44), thus motif features for CTCF and MYC as well as chromatin interaction features are highly ranked. For the remaining 6 transcription factors, we can observe similar results that chromatin features are still the dominative factors, and motif features and chromatin interaction features are the important factors (Supplementary figure 4). However, the feature importance computed by XGBoost just provides a global analysis of feature importance during the training process, and cannot give a deep insight into how each feature independently contributes to predicting CSSBS.

**Figure 6.**
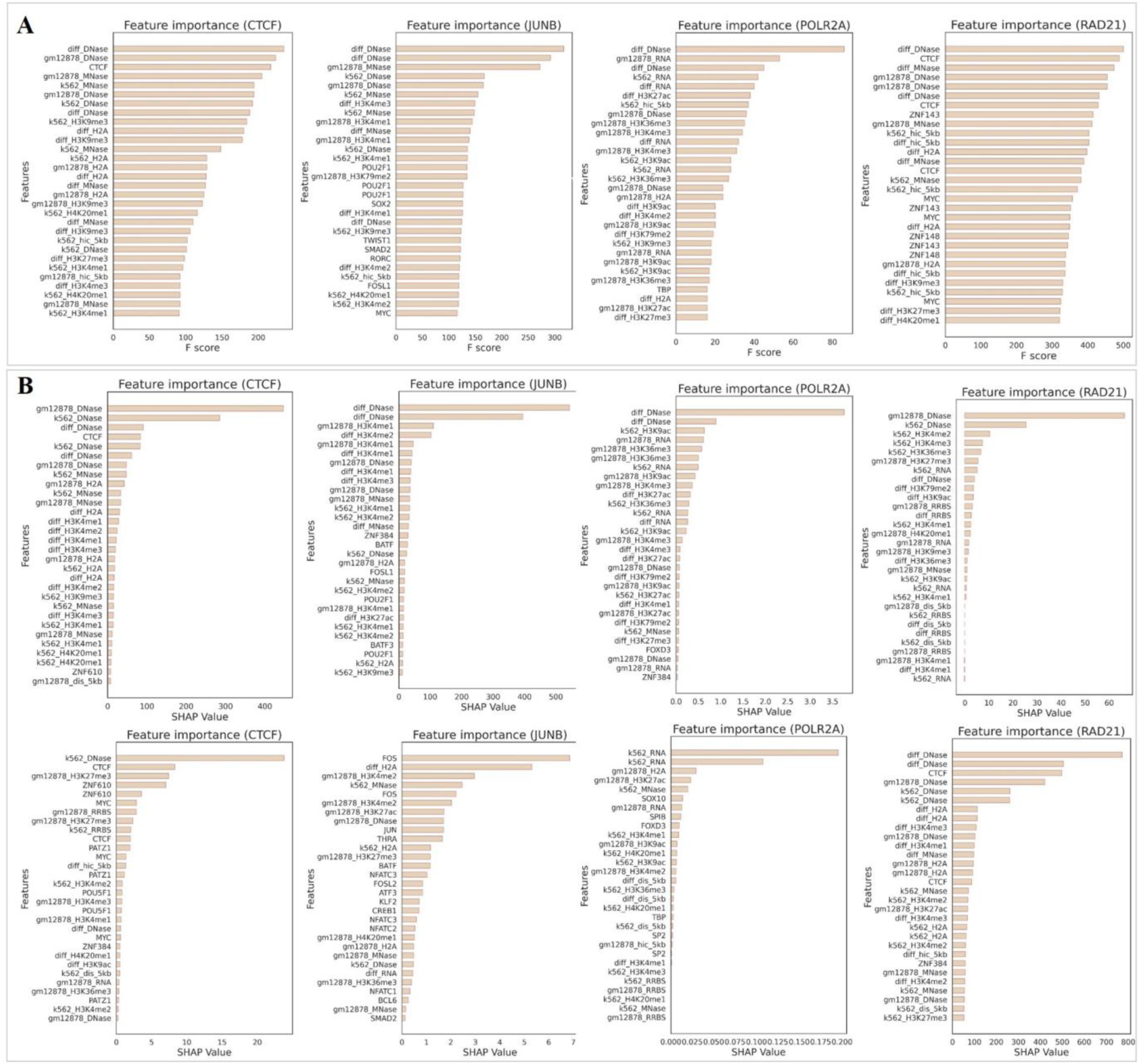
Feature importance analysis to identify independent feature contributions. (A) The global feature importance of the top 30 features across CTCF, JUNB, POLR2A, and RAD21. (B) The top 30 SHAP values of cell-type-specific (top) and shared (bottom) binding sites across CTCF, JUNB, POLR2A, and RAD21. Note that the terms marked by ‘diff’ mean the difference of values between GM12878 and K562.

To alleviate the above problem, we utilized Shapley additive explanations (SHAP) (45) to interpret the outputs of our model and reveal the independent feature’s contribution to predicting CSSBS. Briefly, (i) we separated the test set into cell-type-specific binding sites (positive) and shared binding sites (negative) by labels; (ii) computed the SHAP values of the positive and negative sets respectively, yielding two *n*X*m* matrices where *n* and *m* denote the number of samples and features respectively; (iii) took the sum of each matrix along the sample direction, yielding a vector of length *m*; (iv) took the absolute value of the vector as the final feature importance, representing the independent contribution of each feature to predicting CSSBS; (v) selected the top 30 features with high SHAP values to display. As shown in Figure 6B, we observe that chromatin features, especially DNase-related features, play a main role in the cell-type-specific binding sites while motif features or chromatin interaction features play a main role in the shared binding sites. For CTCF, except itself, other potential cobinding TFs (e.g, ZNF610, MYC) are the main contributors to shared binding sites. For JUNB, DNase features are also the main contributors to cell-type-specific binding sites while its cobinding factor FOS is the most important contributor to shared binding sites. For RAD21, its cobinding factor CTCF plays a very important role in shared binding sites but not in cell-type-specific binding sites. Except for POLR2A, its DNase features are the main contributors to cell-type-specific binding sites while its RNA features are the main contributors to shared binding sites. Similarly, for the remaining 6 TFs, we observe the same results (Supplementary figures 5 and 6). These observations are indeed consistent with the intuition that, cell-type-specific binding sites are characterized by significantly distinct DNase signals while shared binding sites are characterized by similar DNase signals, thus DNase signals are the main discriminator of cell-type-specific binding sites while other features such as cobinding factors or chromatin interactions are the main discriminator of shared binding sites.

### Cross-factor and cross-cell prediction of cell-type-specific and shared binding sites

To evaluate the cross-factor prediction performance of our proposed approaches, we used the model trained on one of the 10 differential binding datasets to test the remaining 9 datasets in rotation. This allows us to study different binding factors in the same cellular environment. The overlapping peaks of different binding datasets can also influence the prediction performance since those overlapping peaks have the same chromatin environment and yield similar feature sets (specifically chromatin features). As shown in Figures 7A and 7B, we not only computed the cross-factor PRAUC of CNN-plus and XGBoost but also the overlapping number between all binding datasets. In general, we find that, in most cases, our proposed approaches perform well for cross-factor prediction of CSSBS, and that CNN-plus is slightly better than XGBoost in terms of PRAUC (0.697 vs. 0.686). For some individual cases, our approaches achieve high PRAUC (0.778 and 0.894) on the MAX and MYC binding datasets in which a large number of peaks between them are overlapped (Figure 7B) since they can form a Myc-Max heterodimer that plays a very important role in cancer cells (46). There is a lot of evidence that cohesin (RAD21) cooperates with CTCF to form CTCF/cohesin anchors and mediate the formation of genome-wide chromatin loops [31, 32], supported by their large number of overlapping peaks (Figure 7B). The models trained on CTCF achieve high PRAUC values (CNN-plus: 0.858, XGBoost: 0.868) on RAD21 while the models trained on RAD21 achieve relative low PRAUC values (CNN-plus: 0.592, XGBoost: 0.741), which demonstrates that RAD21 (cohesion subunit) are not only involved in the formation of CTCF-mediated chromatin loops but also other types of chromatin loops such as promoter-enhance interactions. The model trained on JUNB performs well on POLR2A but poorly on other datasets, suggesting that JUNB can act as a cofactor to engage in the initialization of transcription. Importantly, CEBPB, FOS, and POLR2A rarely overlap with other binding datasets but the models trained on them can accurately predict CSSBS on most of the other binding datasets. This observation implies that there exist some similar patterns of CSSBS between different binding factors in the same cellular environment regardless of distinct sequence specificities between them, demonstrating that our proposed approaches have a good generalization ability of cross-factor CSSBS prediction. To further validate it, we trained a unified CNN-plus model using an integration of the training set from all binding datasets and then used it to measure each test set respectively. As shown in Figure 7C, the unified CNN-plus performs well across all binding datasets, achieving an average AUC and PRAUC of 0.85 and 0.79, further proving the existence of similar patterns of CSSBS between different binding factors in the same cellular environment.

**Figure 7.**
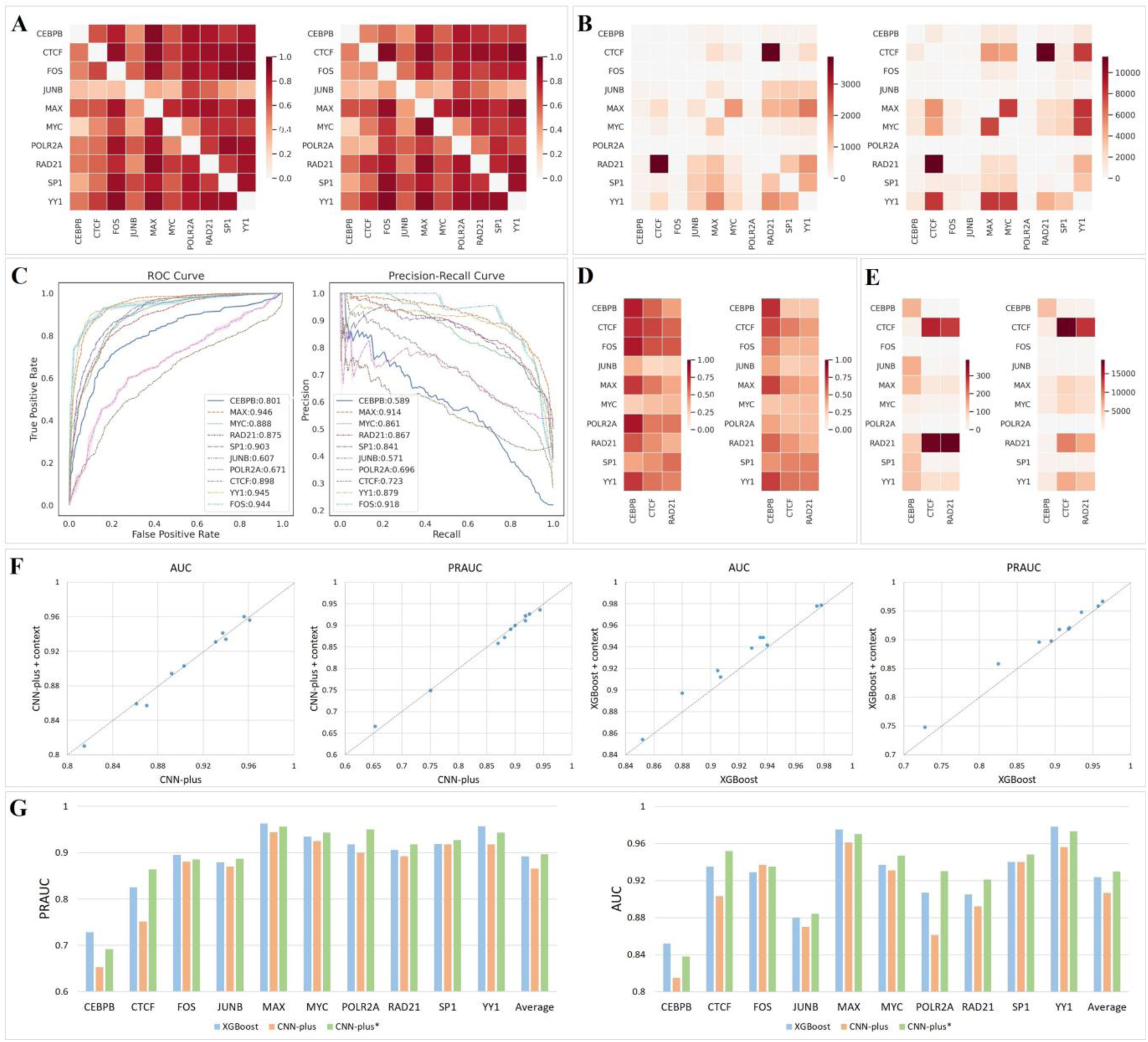
(A) The cross-factor performance of CNN-plus (left) and XGBoost (right) in the same cellular environment, which means that the model trained on one binding factor’s training data from GK was used to test other binding factors’ test data from GK. ‘GK’ denotes GM12878 and K562. (B) The overlapping number of cell-type-specific binding peaks between each pair-wise factor from GK (left) and the overlapping number of shared binding peaks between each pair-wise factor from GK (right). (C) The predictive performance of a unified CNN-plus model on predicting CSSBS, which means that a model trained on all binding factors’ training data was used to measure each binding factor’s test data one by one. (D) The cross-cell performance of CNN-plus (left) and XGBoost (right) in the different cellular environment, which means that the model trained on one binding factor’s training data from GK was used to test three binding factors’ test data from IA. ‘IA’ denotes IMR-90 and A549. (E) The overlapping number of cell-type-specific binding peaks between each pair-wise factor (left) where one is from GK while another is from IA, and the overlapping number of shared binding peaks between each pair-wise factor (right) where one is from GK while another is from IA. (F) The performance comparison of CNN-plus and CNN-plus with additional context information (left), and the performance comparison of XGBoost and XGBoost with additional context information. (G) The performance comparison of XGBoost, CNN-plus, and CNN-plus* where CNN-plus used DNase-related signals while CNN-plus* used all chromatin landscapes.

To evaluate the cross-cell prediction performance of our proposed approaches, (i) we collected additional ChIP-seq datasets of three binding factors including CEBPB, CTCF, and RAD21 from the A549 and IMR-90 (AI) cell lines, and then generated their corresponding differential binding sites by using DESeq2 (see ‘**Differential binding sites preparation**’); (ii) we used the models trained on the 10 binding factors from GM12878 and K562 (GK) to test the three datasets in order. This allows us to study the same binding factor in different cellular environments. Similarly, we also considered the influence of overlapping peaks. As shown in Figure 7D, in general, CNN-plus is much better than XGBoost in terms of PRAUC (0.52 vs. 0.43). In addition, we observe that most of the trained models perform well on CEBPB, demonstrating that this binding factor from AI has the same patterns as the ones from GK. The models trained on CTCF achieve relative low PRAUC values (CNN-plus: 0.673, XGBoost: 0.515) despite a large number of overlapping peaks between CTCF from GK and CTCF from AI (Figure 7E), and XGBoost, heavily dependent on chromatin features, is much worse than CNN-plus, indicating that the chromatin environments around CTCF’s binding peaks in the AI and GK are distinct even if the majority of peaks are at the same genomic position. For RAD21, we observe much lower PRAUC values (CNN-plus: 0.36, XGBoost: 0.35), with a large number of overlapping peaks between RAD21 from GK and RAD21 from AI (Figure 7E), showing that the chromatin environments around RAD21’s binding sites are also distinct. These observations may be caused by the possible reason that there exists structural heterogeneity and functional diversity of topologically associating domains (TAD), associated with CTCF and RAD21 (cohesin subunit), across different cell types in mammalian genomes (47).

### The contextual information of binding sites improves the prediction performance of XGBoost

To explore the effect of the contextual information of binding sites on the performance of predicting CSSBS, we applied CNN-plus and XGBoost to conduct two additional experiments. Briefly, for CNN-plus, we just expanded the original length of peaks from 600bp to 1000bp, since the convolutional layers can automatically learn the contextual information of binding sites in a sliding-window way; whereas for XGBoost, we allowed for the contextual information by expanding the original length of peaks from 600bp to 1000bp, segmenting the peaks of length 1000bp into multiple 200bp genomic bins, and then computing motif features, chromatin features, and chromatin interaction features for each bin. As shown in Figure 7F, CNN-plus with additional contextual information performs worse than CNN-plus, indicating that the contextual information of binding sites is not beneficial to CNN-plus. The result may be caused by the CNN’s intrinsic mechanism that it automatically focuses on the most important features of inputs. However, we find that XGBboost with additional contextual information is better than XGBoost in terms of AUC and PRAUC, showing that the contextual information of binding sites can improve the prediction performance of XGBoost. These observations also suggest that long-range peaks are more suitable for XGBoost than CNN-plus since the inputs to XGBoost are the minimum, mean, and maximum values that characterize the overall distribution of binding peaks.

## DISCUSSION

Unlike TFBS prediction, which is aimed at distinguishing binding sites from non-binding sites, the study of computational prediction of CSSBS is less explored. Although an increasing number of computational methods have been proposed for TFBS prediction and achieved very impressive performance, these methods cannot accurately predict CSSBS when using DNA sequences alone. As our experiments show that LSGKM and CNN, two of the most popular methods, can easily realize the classification of binding and no-binding sites but fail to accurately predict CSSBS. In this paper, thus we proposed two models to predict CSSBS where one is based XGBoost and another is based on CNN. Except for CSSBS prediction (task-B), we constructed other two tasks (task-A: discriminating between GM12878-specific and K562-specific binding sites; task-C: discriminating among GM12878-specific, K562-specific, and shared binding sites.) to verify the prediction performance of our proposed methods. We generated three types of feature sets including motif features, chromatin features, and chromatin interaction features for the XGBoost-based model (XGBoost) while we directly used DNA sequences and DNase-related signals as the inputs to the CNN-based model (CNN-plus). As we expected, our proposed approaches significantly outperform other competing methods on the three tasks. In addition, we explored the importance of different feature sets and identified independent feature contributions by performing ablation experiments and feature importance analysis, observing that chromatin features play a main role in the cell-type-specific binding sites while motif features play a main role in the shared binding sites. Moreover, we investigated the generalization ability of our proposed approaches to different binding factors in the same cellular environment or to the same binding factors in the different cellular environments, finding that the cross-factor and cross-cell prediction performance of CNN-plus is better than that of XGBoost and proving the existence of the similar patterns of CSSBS between different binding factors.

We adjusted DESeq2 to yield binding peaks with significant differences as cell-type-specific (positive) peaks, but it does not mean that these peaks exclusively exist in one cell type. More precisely, a few cell-type-specific peaks exist in two cell types, but the degree of binding in one cell type is much higher than that in another cell type. To compute the difference significance of binding peaks, we just used two biological replicates of binding factors. However, more data, such the cross-platform, cross-project, or cross-lab replicates, should be assembled to acquire more stable difference significance. From the experimental results, the prediction performance of XGBoost is slightly better than that of CNN-plus, perhaps because XGBoost has the benefit of more classification features. To further explain the reason, we re-constructed CNN-plus by encoding DNA sequences and all chromatin landscapes (15 types) together called CNN-plus*and compared it with CNN-plus and XGBoost. As a result, CNN-plus* performs better than CNN-plus on almost all datasets and even better than XGBoost in terms of the average AUC and PRAUC (Figure 7G). Moreover, XGBoost needs much more effort to do feature engineering while CNN-plus only needs DNA sequences and chromatin accessibility signals. The observations suggest the potential advantage of CNN-plus for predicting CSSBS. As for three types of feature sets, motif features are restrictedly extracted from those cobinding factors which have their canonical motifs in the JASPAR database; chromatin features are restrictedly derived from some existing chromatin landscapes such as chromatin accessibility and histone modifications; chromatin interaction features are also restrictedly computed from Hi-C contact maps at single resolution (5kb), but different resolutions (e.g, 1kb, 10kb) may reveal diverse chromatin interaction structures. Even so, we find that the three feature sets are important contributors to predicting CSSBS. These observations lead us to believe that TFs should not be treated as isolated or static, but should be considered in the context of the specific sequence they are binding, their cobinding factors, their chromatin states, and their chromatin spatial structure, which are all likely to play a role in cell-type-specific and shared binding preferences.

As for CNN-plus, (i) we just simply encoded DNA sequences and chromatin accessibility signals, but more appropriate methods for dealing with the inputs need to be further explored, e.g, *k*-mer embedding representing high-order dependencies of nucleotides (48,49), a bimodal neural network for separately handling DNA sequences and chromatin accessibility signals (50); (ii) we just considered a simple CNN architecture composed of three convolutional layers, but advanced DL-based models also need to be further explored, e.g, hybrid neural networks (51,52), transformer architectures (53,54), since they have been proved to be equipped with stronger feature learning ability. Given the massive data being generated by the ENCODE Consortium and other large-scale efforts, there is an excellent opportunity to learn richer representations to more fully understand cell-type-specific and shared binding activities.

