## Supplemental materials for "Computational prediction and characterization of cell-type-specific and shared binding sites"

### Supplementary Figures

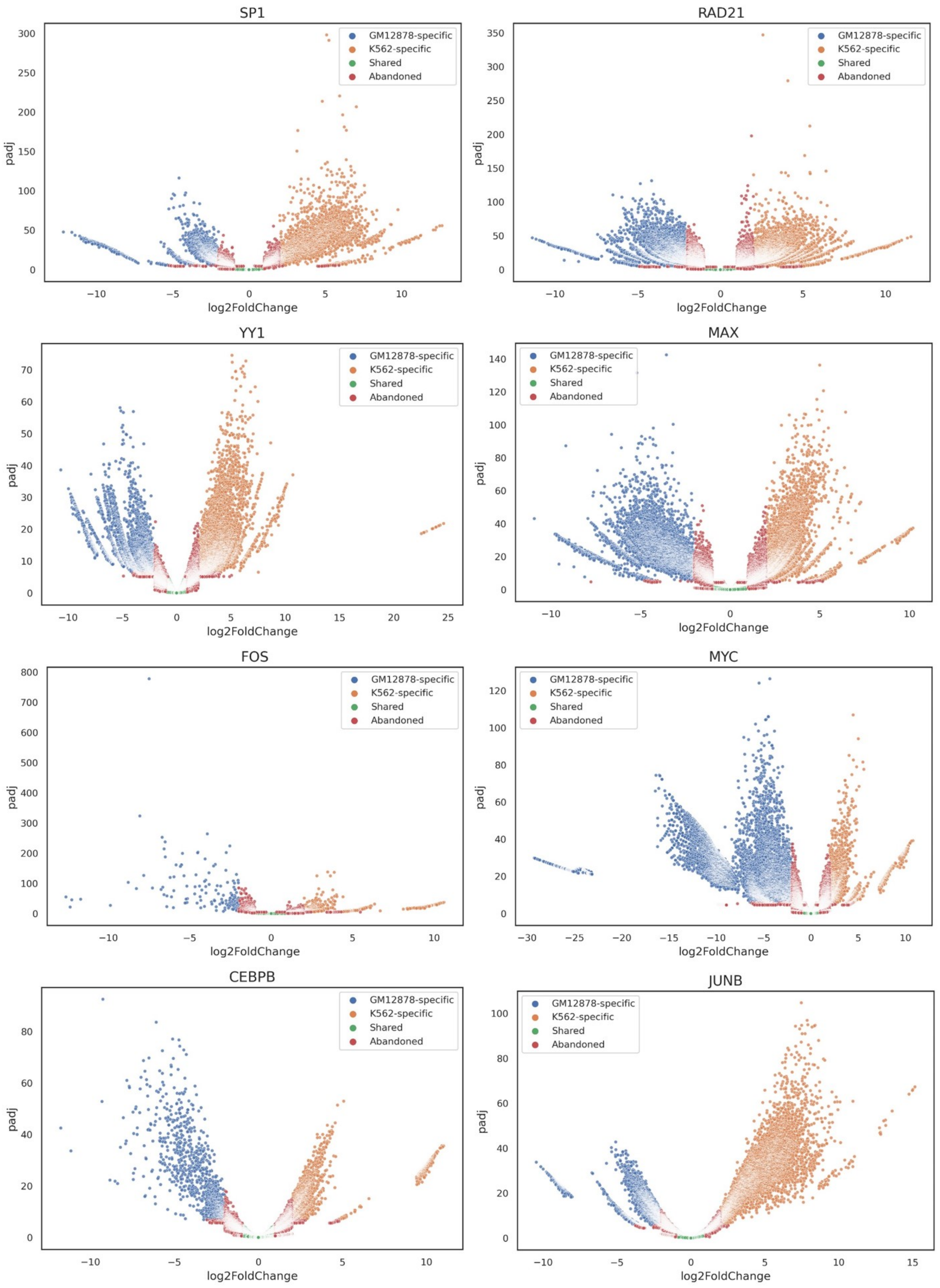


**Supplementary figure 1.** The distribution of binding peaks for SP1, RAD21, CEBPB, YY1, MAX, JUNB, FOS, and MYC, in which GM12878-specific, K562-specific, shared, and abandoned peaks are labelled by different colors.

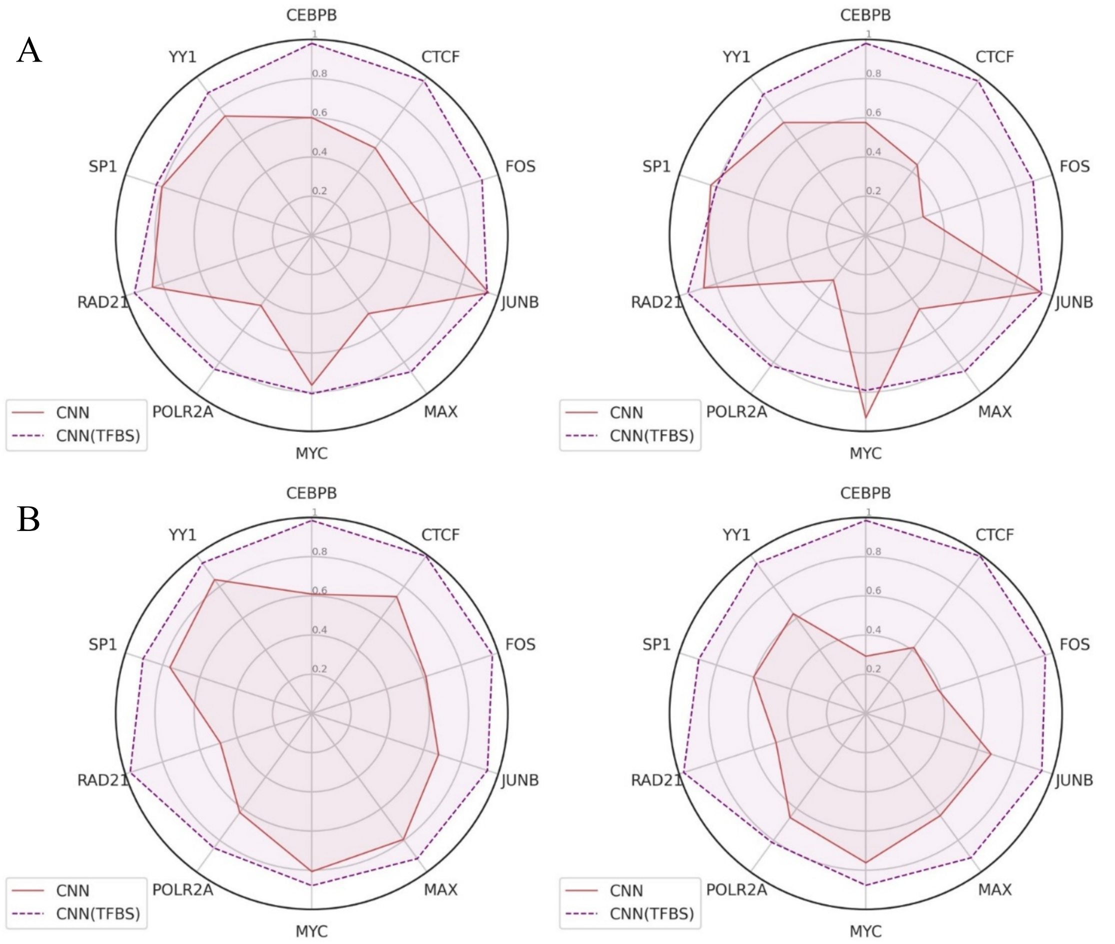


**Supplementary figure 2.** The performance comparison of CNN and CNN (TFBS). (A) The AUC and PRAUC comparison of CNN and CNN (TFBS) across all binding factors where 'CNN' means discriminating between GM12878-specific and K562-specific binding sites while 'CNN(TFBS)' means discriminating between binding sites (GM12878-specific and K562-specific binding peaks) and corresponding non-binding sites. (B) The AUC and PRAUC comparison of CNN and CNN (TFBS) across all binding factors where 'CNN' means discriminating between cell-type-specific and shared binding sites while 'CNN(TFBS)' means discriminating between binding sites (cell-type-specific and shared binding peaks) and corresponding non-binding sites.

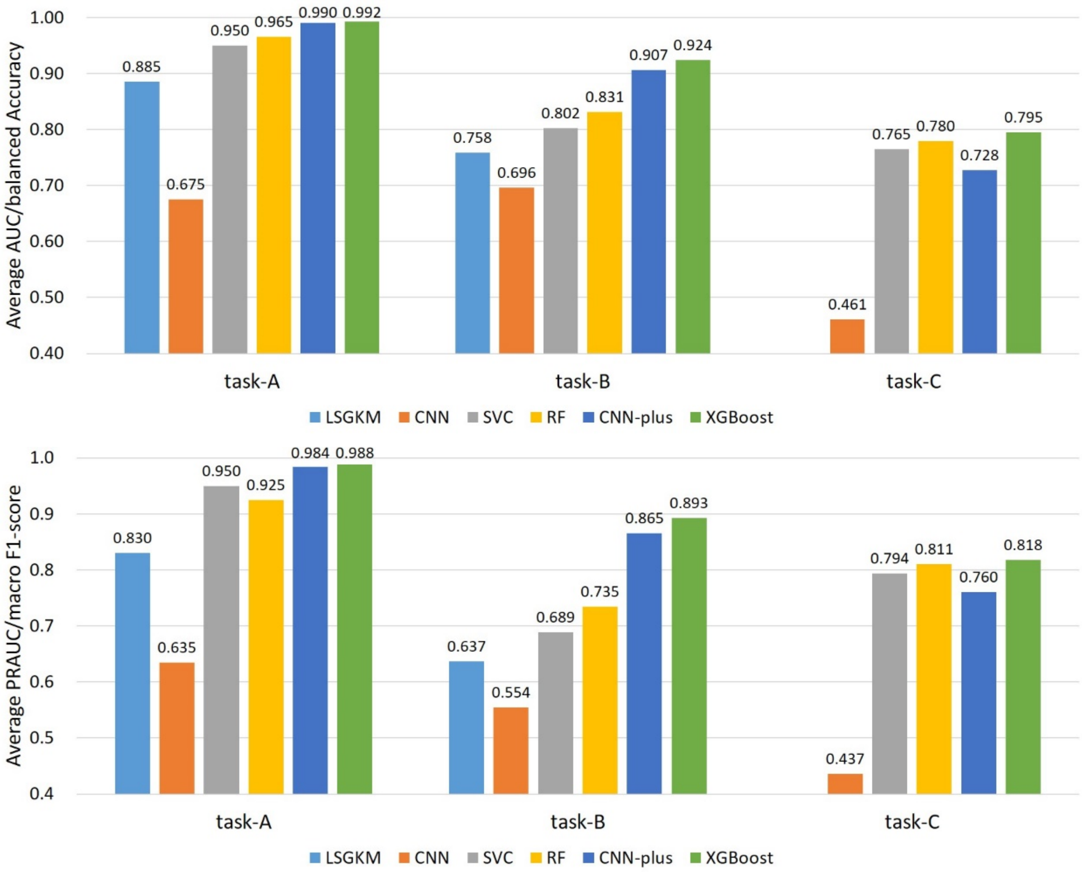


**Supplementary figure 3.** The overall performance comparison of all methods on task-A, task-B, and task-C. For task-A, the average AUC of XGBoost is improved by about 11%, 32%, 4%, 3%, and 0.2% respectively, and the average PRAUC of XGBoost is improved by about 16%, 35%, 6%, 4%, and 0.4% respectively; for task-B, the average AUC of XGBoost is improved by about 17%, 23%, 12%, 9%, and 2% respectively, and the average PRAUC of XGBoost is improved by about 26%, 34%, 20%, 16%, and 3% respectively; for task-C, the average balanced accuracy of XGBoost is improved by about 33%, 3%, 2%, and 7% respectively, and the average macro F1-score of XGBoost is improved by about 38%, 2%, 1%, and 6% respectively.

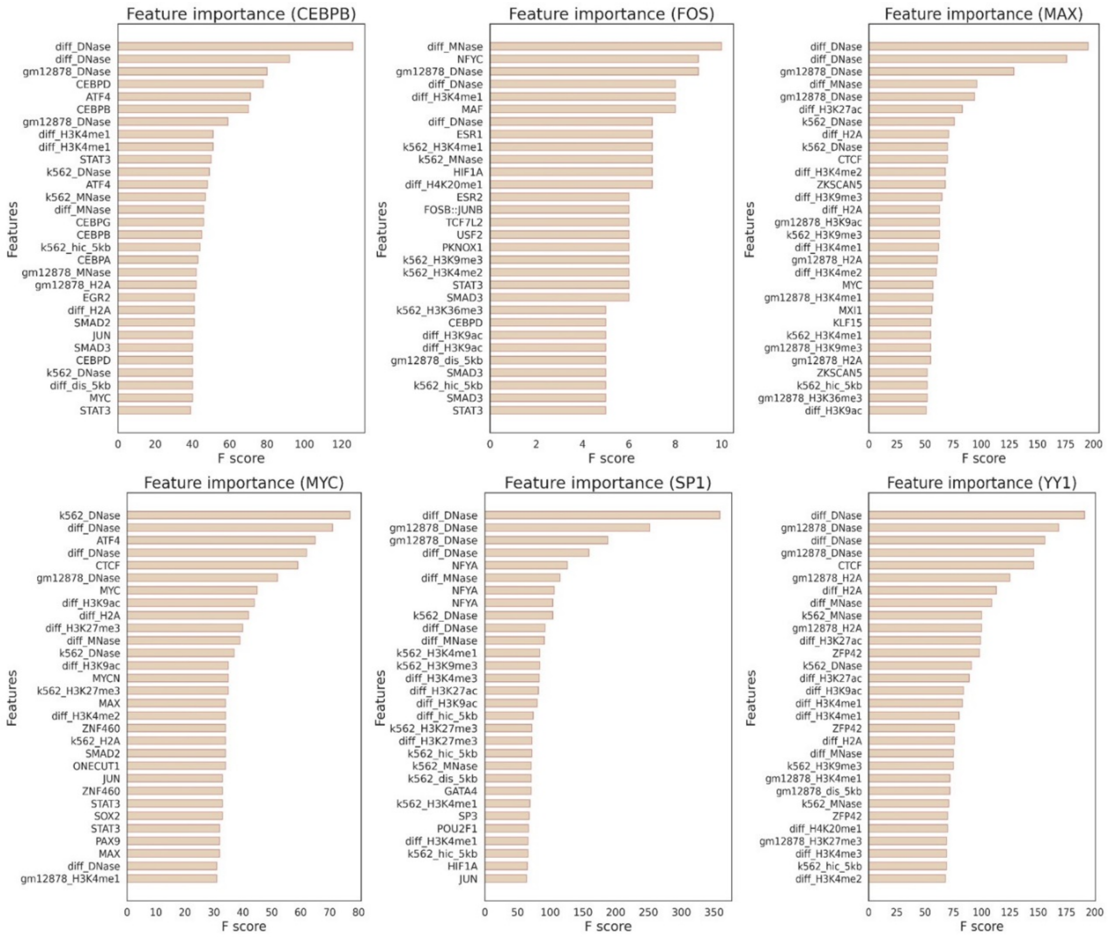


**Supplementary figure 4.** The top 30 feature importance of CEBPB, FOS, MAX, MYC, SP1, and YY1. This type of feature importance provides a global analysis of feature importance during the training process. Note that the terms marked by 'diff' mean the difference of values between GM12878 and K562.

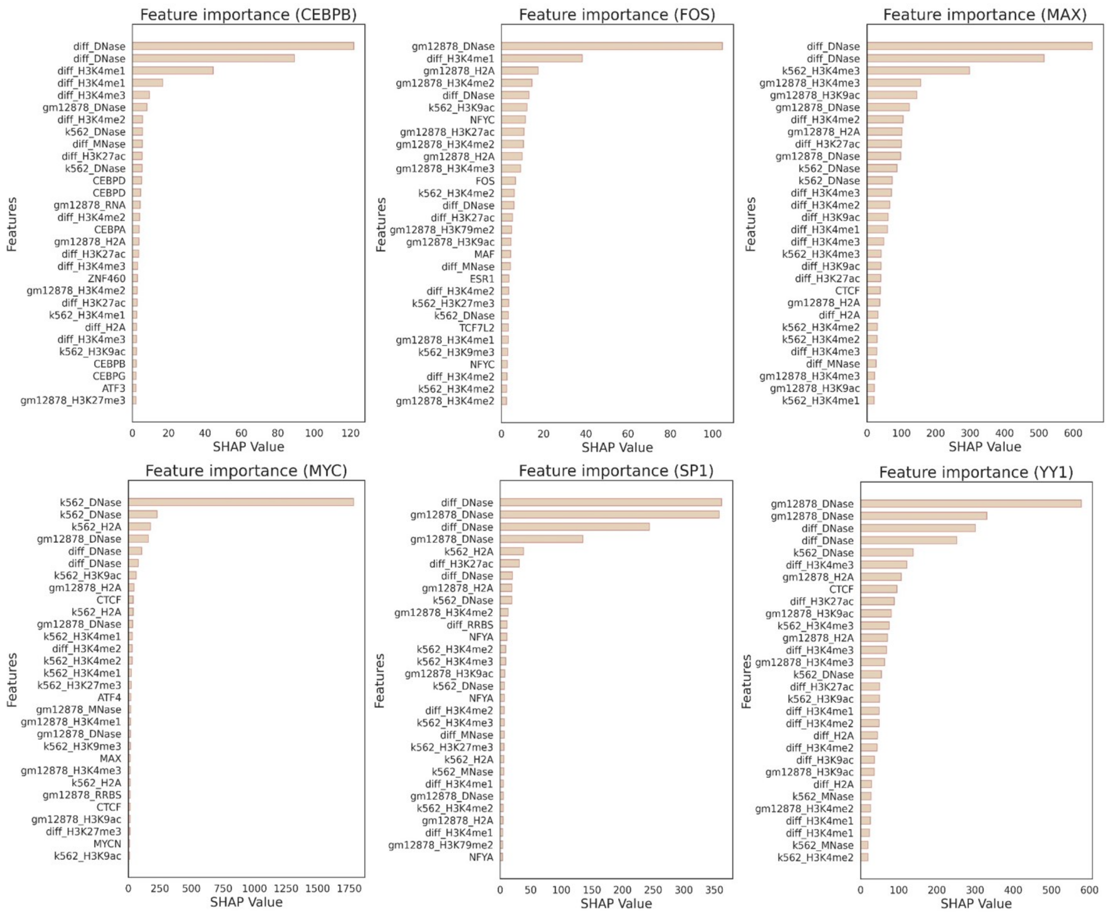


**Supplementary figure 5.** The top 30 SHAP values of cell-type-specific (positive) binding sites across CEBPB, FOS, MAX, MYC, SP1, and YY1. Note that the terms marked by 'diff' mean the difference of values between GM12878 and K562.

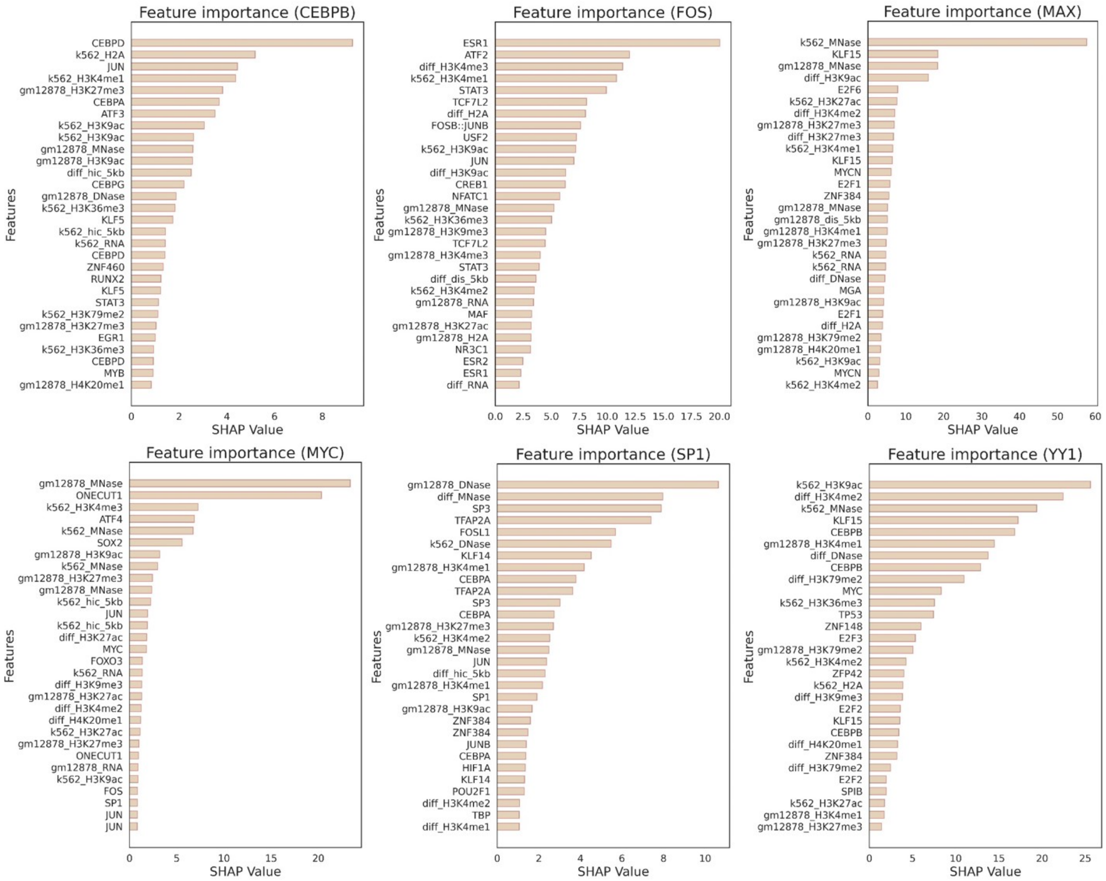


**Supplementary figure 6.** The top 30 SHAP values of shared (negative) binding sites across CEBPB, FOS, MAX, MYC, SP1, and YY1. Note that the terms marked by 'diff' mean the difference of values between GM12878 and K562.

### Supplementary Tables

**Supplementary table 1.** The accession list of ChIP-seq datasets for 10 binding factors from the GM12878 and K562 cell lines. For peaks not provided by ENCODE, we will use the peak caller SPP to generate corresponding peaks.

| GM12878 | Accessions for Peaks | Accessions for Bams |
|---|---|---|
| CEBPB | None | ENCFF573GQJ, ENCFF707LIB |
| CTCF | ENCFF710VEH | ENCFF119SGJ, ENCFF584BRF |
| FOS | ENCFF002COM | ENCFF000VSZ, ENCFF000VTA |
| JUNB | ENCFF939TZS | ENCFF273TMW, ENCFF415ELX |
| MAX | ENCFF083KVY | ENCFF386FSS, ENCFF892REX |
| MYC | ENCFF001USG | ENCFF000ROE, ENCFF000ROF |
| POLR2A | ENCFF120VUT | ENCFF865BUP, ENCFF591BQK |
| RAD21 | None | ENCFF311CJK, ENCFF800DLO |
| SP1 | None | ENCFF000OEL, ENCFF000OEO |
| YY1 | ENCFF967ACD | ENCFF180NKF, ENCFF004ZLO |
| K562 | Accessions for Peaks | Accessions for Bams |
| CEBPB | None | ENCFF059XPJ, ENCFF353LOM |
| CTCF | ENCFF738TKN | ENCFF487UYG, ENCFF496SZR |
| FOS | ENCFF002CVW | ENCFF000YIH, ENCFF000YIG |
| JUNB | ENCFF426DUB | ENCFF095BVD, ENCFF709WYV |
| MAX | ENCFF799HIG | ENCFF938QJA, ENCFF543CKI |
| MYC | ENCFF465JKF | ENCFF058VAU, ENCFF384WMI |
| POLR2A | ENCFF947KPB | ENCFF822HFC, ENCFF359UUS |
| RAD21 | None | ENCFF084HTD, ENCFF330BAK |
| SP1 | None | ENCFF593LCA, ENCFF515QZM |
| YY1 | ENCFF328XKC | ENCFF189XLI, ENCFF872BSD |

**Supplementary table 2.** The accession list of chromatin landscapes from the GM12878 and K562 cell lines.

| GM12878 | URL |
|---|---|
| DNase | https://egg2.wustl.edu/roadmap/data/byFileType/signal/consolidated/macs2signal/pval/E116-DNase.pval.signal.bigwig |
| H2A.Z | https://egg2.wustl.edu/roadmap/data/byFileType/signal/consolidated/macs2signal/pval/E116-H2A.Z.pval.signal.bigwig |
| H3K4me1 | https://egg2.wustl.edu/roadmap/data/byFileType/signal/consolidated/macs2signal/pval/E116-H3K4me1.pval.signal.bigwig |
| H3K4me2 | https://egg2.wustl.edu/roadmap/data/byFileType/signal/consolidated/macs2signal/pval/E116-H3K4me2.pval.signal.bigwig |
| H3K4me3 | https://egg2.wustl.edu/roadmap/data/byFileType/signal/consolidated/macs2signal/pval/E116-H3K4me3.pval.signal.bigwig |
| H3K9ac | https://egg2.wustl.edu/roadmap/data/byFileType/signal/consolidated/macs2signal/pval/E116-H3K9ac.pval.signal.bigwig |

| | |
|---|---|
| H3K9me3 | https://egg2.wustl.edu/roadmap/data/byFileType/signal/consolidated/macs2signal/pval/E116-H3K9me3.pval.signal.bigwig |
| H3K27ac | https://egg2.wustl.edu/roadmap/data/byFileType/signal/consolidated/macs2signal/pval/E116-H3K27ac.pval.signal.bigwig |
| H3K27me3 | https://egg2.wustl.edu/roadmap/data/byFileType/signal/consolidated/macs2signal/pval/E116-H3K27me3.pval.signal.bigwig |
| H3K36me3 | https://egg2.wustl.edu/roadmap/data/byFileType/signal/consolidated/macs2signal/pval/E116-H3K36me3.pval.signal.bigwig |
| H3K79me2 | https://egg2.wustl.edu/roadmap/data/byFileType/signal/consolidated/macs2signal/pval/E116-H3K79me2.pval.signal.bigwig |
| H4K20me1 | https://egg2.wustl.edu/roadmap/data/byFileType/signal/consolidated/macs2signal/pval/E116-H4K20me1.pval.signal.bigwig |
| RRBS | https://egg2.wustl.edu/roadmap/data/byDataType/dnamethylation/RRBS/FractionalMethylation_bigwig/E116_RRBS_FractionalMethylation.bigwig |
| MNase | ENCFF000VME |
| RNA | ENCFF755RBT, ENCFF319UTC, ENCFF975RWM, ENCFF282OWV |
| K562 | URL |
| DNase | https://egg2.wustl.edu/roadmap/data/byFileType/signal/consolidated/macs2signal/pval/E123-DNase.pval.signal.bigwig |
| H2A.Z | https://egg2.wustl.edu/roadmap/data/byFileType/signal/consolidated/macs2signal/pval/E123-H2A.Z.pval.signal.bigwig |
| H3K4me1 | https://egg2.wustl.edu/roadmap/data/byFileType/signal/consolidated/macs2signal/pval/E123-H3K4me1.pval.signal.bigwig |
| H3K4me2 | https://egg2.wustl.edu/roadmap/data/byFileType/signal/consolidated/macs2signal/pval/E123-H3K4me2.pval.signal.bigwig |
| H3K4me3 | https://egg2.wustl.edu/roadmap/data/byFileType/signal/consolidated/macs2signal/pval/E123-H3K4me3.pval.signal.bigwig |
| H3K9ac | https://egg2.wustl.edu/roadmap/data/byFileType/signal/consolidated/macs2signal/pval/E123-H3K9ac.pval.signal.bigwig |
| H3K9me3 | https://egg2.wustl.edu/roadmap/data/byFileType/signal/consolidated/macs2signal/pval/E123-H3K9me3.pval.signal.bigwig |
| H3K27ac | https://egg2.wustl.edu/roadmap/data/byFileType/signal/consolidated/macs2signal/pval/E123-H3K27ac.pval.signal.bigwig |
| H3K27me3 | https://egg2.wustl.edu/roadmap/data/byFileType/signal/consolidated/macs2signal/pval/E123-H3K27me3.pval.signal.bigwig |
| H3K36me3 | https://egg2.wustl.edu/roadmap/data/byFileType/signal/consolidated/macs2signal/pval/E123-H3K36me3.pval.signal.bigwig |
| H3K79me2 | https://egg2.wustl.edu/roadmap/data/byFileType/signal/consolidated/macs2signal/pval/E123-H3K79me2.pval.signal.bigwig |
| H4K20me1 | https://egg2.wustl.edu/roadmap/data/byFileType/signal/consolidated/macs2signal/pval/E123-H4K20me1.pval.signal.bigwig |
| RRBS | https://egg2.wustl.edu/roadmap/data/byDataType/dnamethylation/RRBS/FractionalMethylation_bigwig/E123_RRBS_FractionalMethylation.bigwig |

| | |
|---|---|
| MNase | ENCFF000VNN |
| RNA | ENCFF464HGS, ENCFF295KNU, ENCFF617ZPZ, ENCFF061KDL |

**Supplementary table 3.** The number of GM12878-specific, K562-specific, and shared binding peaks across all datasets.

| Binding Factor | GM12878-specific Peaks | K562-specific Peaks | Shared Peaks |
|---|---|---|---|
| CEBPB | 1109 | 1834 | 11064 |
| CTCF | 3665 | 5693 | 30694 |
| FOS | 143 | 712 | 1680 |
| JUNB | 3847 | 5557 | 10512 |
| MAX | 4684 | 5457 | 16875 |
| MYC | 9439 | 2492 | 16294 |
| POLR2A | 488 | 1556 | 1044 |
| RAD21 | 7474 | 7408 | 15780 |
| SP1 | 2608 | 2677 | 7535 |
| YY1 | 4304 | 5524 | 21568 |
